# Turning point in forest productivity revealed from 40 years of national forest inventory data

**DOI:** 10.1101/2024.04.12.589202

**Authors:** Lionel Hertzog, Jean-Daniel Bontemps, Christian Piedallu, Francois Lebourgeois, Olivier Bouriaud

## Abstract

Aim: Global changes, such as changing climate or disruption in biogeochemical cycles, are affecting forest productivity worldwide. Trends in productivity are depending on the focal spatial scale and on the considered time window, stable trends at large spatial scale can mask divergence at smaller scale while short time windows limit the capacity to reveal non-linear trends such as turning points. Capitalizing on 40 years of national forest inventory data from more than 100 tree species we explored trends in forest productivity at the regional level across 4 biogeographical regions.

Location: France

Time period: 1978-2022

Major taxa studied: 146 tree species

Methods: We fitted two classes of models, a first one explicitly estimating temporal trends and a second one including no temporal components but climatic variables reflecting changing temperature and water availability.

Results: We find a decrease in productivity in 95% of the regions and a high contrast in trend shapes between regions over the period studied: lowland regions with average temperature above 11.9°C showed linear negative trends in productivity since 1985 while colder lowland regions showed hump-shaped trends with turning points between 1985 and 2005, followed by declines in productivity. In mountainous regions, average climate did not appear to be a strong mediator of trend shapes. The temporal trends were reconstituted with high fidelity from the model including only climatic variables implying that changes in temperature and water availability are likely drivers of the reported trends.

Main conclusion: These results illustrate the progression during the last decades of the adverse effects of climate change on forest productivity over the European forests. They suggest the expected changes over the 21^st^ century that will put further pressure on forest productivity, impacting forest carbon sink potential and reducing sustainable rate of timber extraction.

## Introduction

Forests deliver numerous contributions to human societies such as providing timber and non-timber resources but also storing carbon in tree biomass and soils over long timescale (FAO 2022). These contributions are tightly linked to the productive capacities of trees in forests to absorb CO_2_ and turn it into woody biomass. The 20^th^ century witnessed an increase of forest productivity that was reported across countries and biomes in Europe (Spiecker et al. 1996, Boisvenue and Running 2006, McMahon et al. 2010). More recently, evidence is accumulating of declines in productivity for a restricted tree species set (Charru et al. 2017, Martinez del Castillo et al. 2022, Mäkkinen et al. 2022), suggesting that a turning point may have been reached to an extent that remains unspecified to date. Widespread growth declines and ultimately increased mortality would have critical implications in the context of forest adaptation to climate change (Jandl et al. 2019), and also climate change mitigation through carbon sequestration (IPCC 2022).

Forest productivity is controlled by numerous factors and constraints including climatic factors such as temperature and water availability. Climate change is changing the distribution of these factors, mostly in the sense of increasing constraints and affecting forest productivity in various ways. For instance, warmer temperatures over the last decades were reported to increase the productivity of temperate forests worldwide (Boisvenue and Running 2006) through higher metabolic rates (Delpierre et al. 2009) and contributed to a lengthening of the growing season (Myneni M et al. 1997). Longer growing seasons lead to earlier bud bursts for numerous species across Europe (Menzel et al. 2006) stimulating productivity but also increasing the risk of bud’s mortality in spring due to late frosts (Zohner et al. 2020) or to nutrient loss in autumn if deciduous trees are not able to resorb leaf nutrients before leaf fall (Estiarte and Peñuelas 2014). Saxe et al. (2001) reported in their review that numerous temperate and boreal tree species responded positively to increasing temperature in the past decades. However, a continuing increase in temperature will also gradually increase the constraint on water availability for forest productivity in climate context that were up until now mainly temperature limited (Choat et al. 2012, Ols et al. 2020, Babst et al. 2019). Climate change impacts on the precipitation regime is less clear and more variable regionally, with mainly an increase in precipitation amount for middle and high latitudes and a decrease near the tropics, but also with significant changes in seasonal distribution, with longer and more frequent droughts followed by heavier rainfall events (IPCC 2022). Recently, repeating summer heat waves combined with summer drought led to declining ecosystem productivity in forests (Scharnweber et al. 2020, Matula et al. 2023, but see Salomon et al. 2022). Ultimately the response of forest productivity to changes in temperature and precipitation regimes will depend on whether and how these factors are limiting tree growth. Trends in forest productivity are therefore variable through space at the regional scale with potential divergence between regions positively or neutrally affected by the recent climatic changes and other regions negatively impacted (Charru et al. 2017) but also potentially regions showing non-linear trends with turning points. Future trends are uncertain, although increased frequency and severity of extreme climatic events in interaction with biotic stressors will most probably lead to productivity declines in forest and natural ecosystems in general (Morales et al. 2007, Zhang et al. 2022).

Studies on forest productivity trends and drivers mostly focus on a limited number of tree species within defined regions in simple stand structure and composition (see Boisvenue and Running 2006 or Kahle et al. 2008 for reviews) in order to explicitly control for confounding factors in observational settings and to pinpoint trends to given tree species, usually of economic interest or with widespread distributions. While this approach has been successful in revealing environmental- and climate-driven trends (Charru et al. 2017, Ols et al. 2020, Pretzsch et al. 2023) it lacks systemasticity regarding the tree species, stands and regions considered (but see Henttonen et al. 2017). Furthermore, this approach of focusing on a limited set of tree species-regions couples limits the climatic gradients considered, potentially reducing the power of detecting climatic drivers behind the trends identified. At the other extreme, large-scale studies usually do not assess regional trend divergence with few exceptions such as Wang et al. 2023 in Canada or Pretzsch et al. 2023 in Europe. In the context of adapting forest management regimes to climate change, policy-makers and forest managers need systematic assessment of forest productivity trends and their drivers for a large number of species and over broad ecological gradients. Such systematic assessment can be achieved using forest inventory data (see for instance Henttonen et al. 2017). Regional-scale studies of productivity trends across tree species using inventory data requires stability in tree species composition within the considered temporal window. Indeed, given the large difference in productivity between species, shifts in species composition can be a confusing factor of trends in forest productivity. Over long-time scales, such as centuries, tree species composition shifts, especially in the context of climate change (Davis 1983). At medium time scales, such as decades, the evidence of tree species composition shifts is thinner, for instance, using an inventory of more than 6000s stands in North America; Boisvert-Marsh et al. (2014) found no significant latitudinal shifts in adult tree distribution in 10 out of 11 studied species. The review of Lenoir et al. (2020) based on 325 studies also found no clear signal of latitudinal distributional shifts for tree families. The current evidence is, therefore, that tree species composition was relatively stable in the recent decades at the regional scale, consistent with Loelhe and LeBlanc 1996, opening the option of systematic assessment of temporal trend in forest productivity at the regional level from inventory data.

In order to assess the reality and extent of a recent turning point in European forest productivity we considered Western Europe as a case study, using the French National Forest Inventory (NFI). French forests are distributed across various climatic zones and represent almost all major European forest types (Leguédois et al. 2011), from dry Mediterranean systems to cold semi-continental with several mid- and high-altitude mountain ranges, making France a representative case study of a broad number of European forests. We aimed at identifying potential climatic drivers of the trends to inform forest policy and management adaptation to climate change. Given the current knowledge gap associated to a lack of systematic information, our approach embraced all tree species and stand structures present at the regional scale, that we defined here as homogeneous subnational geographical units, to allow a macro-screening of trend variability. To achieve this, we used data from the French NFI based on field measurement on around 300’000 field plots totalling around 4 Mio sampled trees from more than 100 species between 1978 and 2022. The French NFI is a systematic and representative survey of forests over the national (metropolitan) territory which provides unbiased estimation of numerous forests attributes, including productivity. The French metropolitan territory encompasses, according to the European Environmental Agency classification, 4 biogeographical regions that together cover 75% of the European Union land area (Roekaerts, 2002). Previous works have shown the usefulness of the French NFI data to study climate – forest productivity relationship (for instance Charru et al. 2017 or Ols et al. 2021). Our main objectives were: (i) reveal and contrast recent trends of forest productivity at the regional level across 4 biogeographical regions, with the aim to identify a potential turning point in productivity, and (ii) identify potential climatic drivers of these trends.

## Material and Methods

### French National Forest Inventory data

The French National Forest Inventory (NFI) was established in 1958 and digitalized records are available from 1978 onwards. Between 1978 and 2004 the NFI was conducted at county level (NUTS 3 level of the EU system) with inventories approximately every 12 years within each county. Starting in 2005, the NFI is conducted countrywide every year. The sampling method of the French NFI is based on a nationwide grid tesselating the territory onto equally-sized cells in which points are randomly drawn to form the sample of field plots. The French NFI belongs to the family of the two-phase sampling surveys combining a first-phase sample for vegetation and land-use classification, and a smaller nested second phase sample for field measurements (Bouriaud et al. 2023). The number of field plots varied between the years, before 2004 there were around 1200 plots per sampled county and after 2005 around 7000 plots nationwide. In total, data on 345’820 plots are available for computation. From the sampled data, unbiased mean and precision estimates of forest attributes are computed using a two-stage sampling for post-stratification estimator. Further descriptions of the French NFI sampling and estimation methods are given in Bouriaud 2020 and Bouriaud et al. 2023. The French NFI allows the estimation of forest attributes for different spatial domains including *forest regions*. Forest regions form a spatial partition of the territory, aimed at representing homogenous abiotic and forest conditions. 86 forest regions are defined in France with an average area around 6400 km² (Table S1). These forest regions are nested within 11 larger biogeographical regions representing broad climatic and topographic gradients (Fig. 1). Three biogeographical regions fall within the oceanic range (named: Grand ouest, Centre nord and Sud ouest), one within a semi-continental climate (Grand est), three are mid-altitude mountain ranges (Vosges, Jura and Massif central), two are high-altitude mountain ranges (Alpes and Pyrénées) and two have a Mediterranean climate (Méditerranée and Corse). Detailed information on the biogeographical and forest regions is available from IFN (2011) and Cavaignac et al. 2004 and the list of dominant tree species per forest region is given in Table S1. With the EU bioeconomy strategy increasing attention has been paid on regional forest resources, a scale at which systematic knowledge is currently missing.

**Figure 1:**
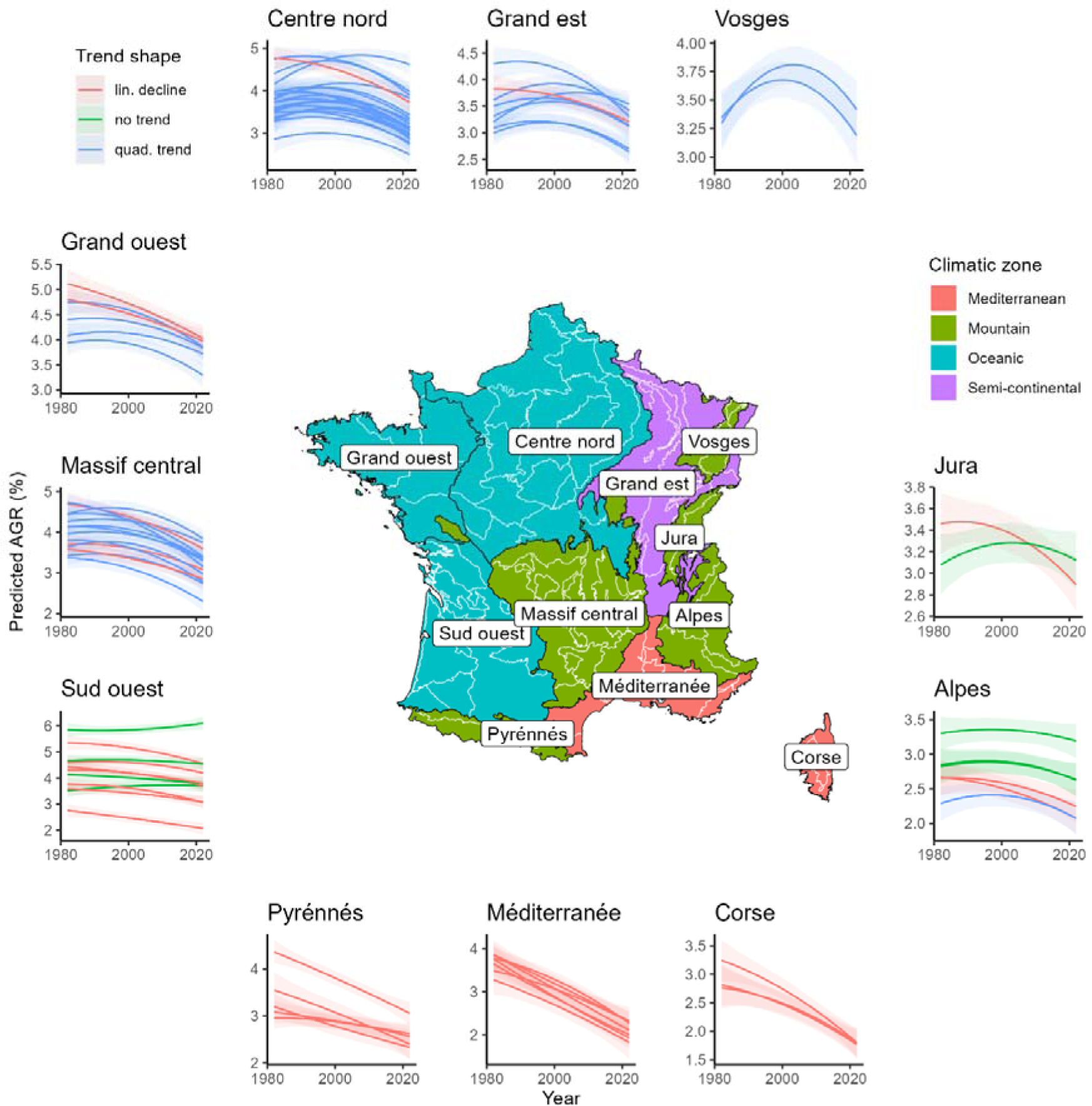
Centre: Map of the 11 biogeographical regions with names and black contours together with the 86 forest regions embedded within with white contours. The filling colors represent the climatic zones associated to the biogeogrpahical regions. Outside: trend in absolute growth rate predicted by the model defined in equations 1-4 at the level of the forest regions. The blue lines indicate substantial evidence of a turning point (P(β_2_ < 0) > 0.9) while the red lines the converse. Green lines indicate forest region with no substantial evidence for temporal trend. The envelopes are the 90% credible intervals around the predicted trends.

### Stocks, productivity and stand development data

Field measurements include number of trees, circumference at breast height and, on a subset of trees tree cores. In total around 4Mio trees were measured in the field over the period. These variables are measured within circular plots of sizes depending on the diameter classes of the trees (for instance trees of a diameter at breast height between 7.5 and 22.5 cm are measured in a 6 m radius plot). Trees with diameter at breast height smaller than 7.5 cm are not measured and are not taken into account in metric computation. Each tree is given a weight depending on plot size to scale the estimation per hectare of forest. We used here the aboveground merchantable stem volume of the living trees (up to a minimum diameter of 7.5 cm) as a proxy for forest growing stock. Volume is estimated per tree from the circumference at breast height and total height using tree species-specific allometric equations described in Morneau 2018. These estimations are weighted and summed per plot to provide a volume in m^3^ per hectare.

For productivity we used the estimated tree-level annual growth in merchantable stem volume (m^3^ / ha / year) computed from tree cores taken at breast height on at least one tree per species and per diameter class that quantify the length of radial growth from the last 5 fully formed tree rings. This measure allows an estimation of tree volume 5 years ago which is then used to compute the annual growth in volume per tree and then per hectare of forests (Ols et al. 2020). Total merchantable volume of the newly recruited trees crossing the accountability threshold of 7.5 cm of diameter at breast height over the last 5 years are added to the estimated productivity for their total volume. The estimations of stocks and productivity were based on living trees and did not consider the production and the volume of dead and harvested trees (i.e net productivity). Estimation of gross productivity taking into account the productivity from harvested trees and of volume from harvested and dead trees are available only for the years 2010 until 2017 based on the recent resurvey approach implemented after 5 years. Over that period, a gross rate of productivity was therefore computed and compared with the net rate, showing a very tight correlation (see Fig. S1), and justifying the use of net productivity as a proxy for forest productivity. Growing stocks have markedly increased in French forests over the last 40 years (IGN, 2023), given that productivity is monotonously linked to growing stocks and to control for this confounding effect, the absolute growth rate (AGR, productivity divided by growing stocks) was computed. Finally, stand development stage was measured by the quadratic mean diameter of forest plots. Growing stock, productivity and stand development stage were estimated at the level of the forest regions and for non-overlapping temporal windows of 5 years, so 9 temporal replicates per forest regions were available over the considered period. This 5-year temporal window is routinely used for the reporting of the official forest statistics from the French NFI in order to increase the precision of the estimate. Before 2004 forest inventory was conducted at NUTS-3 level. Therefore, several separate inventories were potentially sampling the same forest regions within a given 5-year temporal window. These separate estimations were aggregated using a weighted mean with the number of field sampled points as weights. After 2005 the estimation algorithm allows direct computation of the estimates at the level of the forest regions for any temporal window. Estimations based on low numbers of field plot (less than 100 for estimations before 2005 and less than 30 for estimation after 2005) were dropped from the analysis, as a result one forest region could not be analysed so 85 were used in subsequent analysis.

Data gathering and manipulation operations were conducted in R v4.3 (R Core Team 2023) using the packages tidyverse v2.0 (Wickham et al. 2019), data.table v1.14 (Dowle and Srinivasan 2023) and inventR v4.3 (Morneau 2023).

### Climatic data

We used the dataset developed in Piedallu et al. (2016). This dataset is derived from homogenised meteorological station records mapped on a 1km grid using empirical and process-based models as well as interpolation of model residuals (Ninyerola et al. 2000). The variables gathered were the seasonal mean, minimum and maximum temperature as well as seasonal soil water deficit (difference between potential and actual evapotranspiration) computed from the soil water balance estimated from Thorthwaite’s method (Piedallu et al. 2013). Data were gathered between the year 1978 and 2020. For each variable and each pixel, a climate normal was derived as the 30-year average between 1980 and 2009, then the absolute differences (anomalies) between each annual value and the climate normal were computed. These anomalies were then averaged per forest region and 5-year temporal window to match the productivity data. The 12 temperature variables (4 seasons x 3 indicators) were highly correlated, a PCA was run and the unit observation scores on the first axis, representing 73% of the total variation, were kept for further analysis. All 12 temperature variables were correlated to this first axis that represents the main axis of variation in temperature anomalies across seasons (see Table S2 for the variable loadings). The anomalies in water deficit for the growing season (spring and summer) were also kept. Correlation between these select variables (first PCA axis, water deficit in spring and summer) ranged from 0.39 to 0.64. The relative anomalies (anomalies / climate normal) were also derived, but the results obtained were extremely similar to the absolute differences. We therefore only present results based on the absolute differences. Manipulation of the climatic data was performed in R using the packages terra v1.7 (Hijmans 2023) and tidyverse.

### Productivity Trend analysis

To answer our study objectives, hierarchical linear regression models were developed, with forest regions within a given 5-year temporal window considered as observation units. Absolute growth rate (AGR) was the response variable. Independent variables included the quadratic mean diameter (QMD) to control for stand maturity, and *year* terms to represent temporal trends in the AGR. All covariates in the models were standardized to unit variance and centred to their mean prior to the fitting. To test for the shape of the temporal trends, and namely for the existence of a turning point (maximum), orthogonal polynomials of *year* from degree 1 to 3 were used in three nested models. Higher polynomial degrees were not considered to avoid overfitting, considering that a maximum of 9 temporal points were available. The fitted models were of the form:

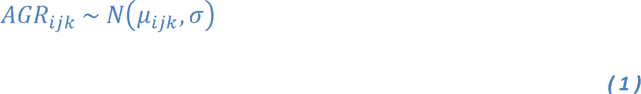

Where AGR is the ratio of productivity over stock, i is an index for the biogeographical regions (from 1 to 11), j is an index for the forest regions within the biogeographical regions (from 1 to 85) and k an index for the unit observation (the sample size, from 1 to 632), N is a Gaussian distribution with mean µ and standard deviation σ.

The mean is conditional to model covariates, and for the polynomial of degree 1 yields:

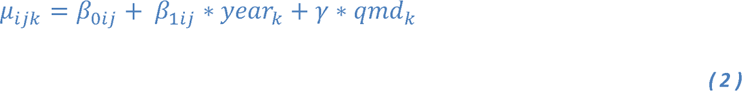

The hierarchical structure in the data (forest regions within larger biogeographical regions) was considered by allowing the intercept and the polynomial parameters of year to vary at both levels formed by biogeographical and forest regions, using a nested structure. Thus, the parameters β are regression intercepts and the year slopes varying between biogeographical regions (i) and between forest regions within biogeographical regions (j). The models for the polynomial degree 2 and 3 have additional quadratic (β_2ij_) and cubic terms (β_3ij_) with the same hierarchical structure.

The regression coefficient β_*ij_ are defined as follow:

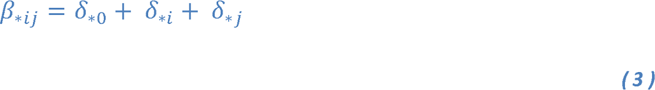

With the δ_*i_ and δ_*j_ sampled as follow:

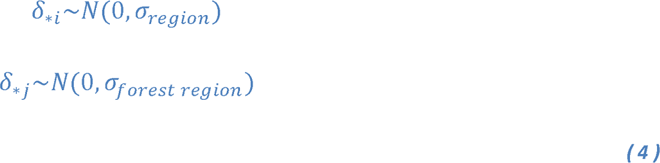

These terms allow for deviation in the regression coefficient between the biogeographical regions and the forest regions within the biogeographical regions. The hierarchical structure also allows for partial pooling of the estimated coefficients across forest regions, which provide an efficient compromise between complete pooling (one coefficient across all regions) and no pooling (all regions get independent coefficients, Gelman and Hill 2006). The models were fitted using the library brms v2.20 (Bürkner 2018) from R that fits statistical models within a Bayesian framework using the Stan language as backend. Default priors (see Table S3) and sampling settings were used: 4 chains were run for 2000 iterations with the first 1000 iterations used as burn-in for the algorithm. Convergence of model parameters was evaluated using Rhat coefficients, a threshold of 1.01 was used following Vehtari et al. (2021). Model fitness was evaluated using residual plots from posterior predictive distributions. The following plots were made: residuals vs fitted values (check for homoscedasticity), residuals vs model covariates (no deviation from linear assumptions) and posterior predictive distribution against the distribution of the observations (predictive ability of the model). To assess the polynomial order best fitting the data, expected log-pointwise density were derived using leave-one-out cross validation as implemented in the package brms which quantifies the leave-one-out information criterion that is used to compare models (Vehtari et al 2017). This criterion measures the relative degree of adequacy of the model to the data, smaller values indicating better fit.

### Climate change analysis

To further understand the relationship between climate change and trends in the rates of productivity at a regional level, we built a set of hierarchical models with a structure similar to the model defined in equations 1-4. The following covariates were tested in nested models: (i) the position of forest regions within a given temporal window on the first PCA axis of the 12 temperature variables (pc1), (ii) the squared position on the first PCA axis to account for potential hump-shaped response of productivity to warming (pc1²), (iii) the summer water deficit (wd_summer) and (iv) the spring water deficit (wd_spring). First, the following full model was built:

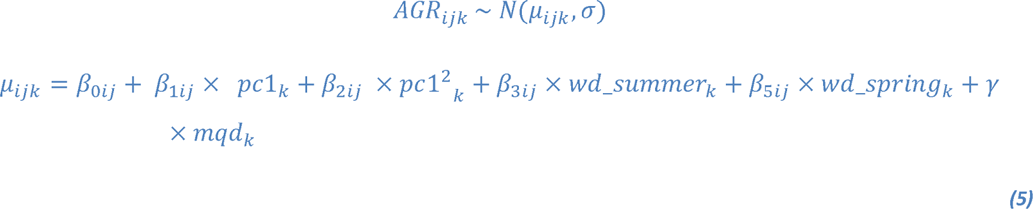

The regression coefficient β_*ij_ being sampled in the same way as in the trend model defined in equations 1-4. The model was fitted using brms with the same settings as presented above. From the full model, the regression coefficients at the level of the biogeographical regions were extracted to explore variations in response to climate change. In addition to the full model, the following reduced models were built: (i) pc1, (ii) wd_summer + wd_spring, (iii) pc1 + wd_summer + wd_spring, (iv) pc1 + pc1². This stepwise approach was used to assess how closely the climatic models could reproduce the temporal trends but also to identify the most parsimonious set of covariates that best correlated with the fitted temporal trend. The predicted AGR for the different temporal windows and forest regions were predicted from the five climatic models and correlated to the fitted temporal trends.

## Results

### Data exploration

The merchantable stem volume per hectare, the chosen proxy for the growing stock density, varied spatially between 21 and 344 m^3^/ha with a mean of 155 m^3^/ha between 1978 and 2022. The forest regions with the largest stock densities were in the Vosges and Jura mid-elevation mountains, and the lowest stock densities were observed in the Méditerranée and Corse Forest regions. Stock density has been continuously increasing between 1978 and 2022 from 119 to 180 m^3^/ha on average. This increase was observed in most forest regions (Fig. S2). Stem volume growth ranged from 0.7 to 10 m^3^/ha/year with a mean of 5 m^3^/ha/year. The highest average productivities over the studied period were also found in the forest regions of the Vosges Mountains and the lowest were found in Méditerranée and Corse Forest regions. Overall, productivity increased from 1980 to 2012 but declined afterwards. At the regional level, this trend was visible in forest regions of Northern France but absent from forest regions situated in southern France which showed more stable trends in productivity (Fig. S3). Absolute growth rate, the ratio between productivity and growing stocks, ranged from 2.2 to 4.9% with an average of 3.5%. Greatest absolute growth rates were found in the forest regions of the oceanic Grand Ouest and the lowest in the forest regions of Corse. Absolute growth rate declined in all biogeographical regions over the study period, the largest declines were found in the Méditerranée forest regions and the lowest declines in Corse (Table 1). Quadratic mean diameters ranged from 0.38 to 0.98 m with an average of 0.61 m. The forest regions with the greatest developmental stages were in the Vosges mountains and the forest regions with the most restricted developmental stages were in the Méditerranée. Quadratic mean diameter increased over the studied period from 0.53 m in 1978 to 0.66 m in 2022, this increase is apparent in most forest regions (Fig. S4).

**Table 1:** Absolute growth rate (AGR in %) at the beginning (1978-1982) and the end (2020-2022) of the considered temporal window per biogeographical regions and for the entire country as estimated from the French National Forest Inventory. The climate zone corresponding to the biogeographical regions is given as well as the absolute difference between the maximum and minimum absolute growth rate values observed. The posterior probability that the AGR in 2020-2022 is lower than in 1978-1982 based on the model defined in equation 1-4 is also given. A map of the biogeographical regions is given in Figure 1.

| Biogeographical region | Climate | AGR 1978-1982 | AGR 2020-2022 | Absolute difference max – min | Prob. AGR 2022 < AGR 1982 |
| --- | --- | --- | --- | --- | --- |
| Grand ouest | Oceanic | 4.86 | 3.44 | 1.44 | 0,99 |
| Centre nord | Oceanic | 4.26 | 2.90 | 1.36 | 1 |
| Grand est | Semi-continental | 3.37 | 2.88 | 0.90 | 0,97 |
| Vosges | Mountain | 2.82 | 2.78 | 0.78 | 0,75 |
| Jura | Mountain | 3.33 | 2.57 | 0.75 | 0,89 |
| Sud ouest | Oceanic | 4.72 | 3.90 | 1.03 | 0,99 |
| Massif central | Mountain | 4.30 | 3.01 | 1.31 | 1 |
| Alpes | Mountain | 2.79 | 2.22 | 0.57 | 0,95 |
| Pyrénées | Mountain | 3.74 | 2.49 | 1.25 | 1 |
| Méditerranée | Mediterranean | 4.65 | 2.67 | 1.98 | 1 |
| Corse | Mediterranean | 2.61 | 2.16 | 0.47 | 1 |
| Entire France |  | 3.91 | 2.90 | 1.01 | 0,99 |

### Model selection and overall model fitness

The model defined in equations 1-4 was fitted with three different forms of temporal trend using polynomials: linear, quadratic and cubic. The comparison of the trend shapes revealed that the data supported, overall, temporal trends of higher complexity than linear trends (Table 2). The difference in the expected log-pointwise density between the quadratic (polynomial of degree 2) and the cubic (polynomial of degree 3) trend was small and within the estimated residual error. The simplest model (quadratic temporal trend) was therefore retained, for all forest regions, for further analysis. The selected model showed convergence on all parameters and the residual plots showed no significant deviation of the residual pattern from the expectation (Fig. S5, S6). The R² of the model taking into account all parameters (including hierarchical terms, equations 1-4) was 88%. In contrast, the R² taking into account only population-level parameters (no hierarchical terms included, equations 1-2) was 28%. Thus, regional variations in average absolute growth rate and in temporal trends were responsible for the major part of the signal (60%). Increasing quadratic mean diameter led to decline in absolute growth rate of -0.34 (95% CrI -0.41 - -0.28) for a unit (0.09m) increase in quadratic mean diameter. The temporal trend in absolute growth rate showed a moderate increase between 1978 and 1988 from 3.54 (95% CrI 3.07 – 3.99) to 3.62 (95% CrI 3.21 – 4.02), followed by a decrease reaching 2.96 (95% CrI 2.52 – 3.38) in 2022.

**Table 2:** Comparison in the expected log-pointwise density between the models with different polynomial degrees. Negative values indicate support for the first model in the comparison and positive values indicate stronger support for the second model.

| Comparison | Expected log-pointwise density difference (with s.e.) |
| --- | --- |
| Linear vs quadratic | 57 ± 11 |
| Quadratic vs cubic | -4 ± 5 |
| Linear vs cubic | 61 ± 13 |

### Region-level trends in AGR

The regression coefficient of the temporal trend (β_1_ and β_2_ in equation 2) showed large variations between the forest regions (Table S4). The model was therefore able to capture the large variations of the trends in absolute growth rate between the forest regions, and this variation was mainly associated to the linear coefficient (β_1_) of the trend. The predicted trends at the level of the forest regions showed variations in their trajectory with around 50% (45) of the forest regions showing clear hump-shaped trends (P(β_2iJ_ < 0) > 0.9,Fig. 1). In these forest regions, the hump reveals the existence of a turning point in absolute growth rate, and suggested an inversion of the trend over the period under study. The turning point was on average located in 1994, with the earliest optimum being in 1982 (or earlier) in two forest regions of the Massif Central and the latest in 2007 in two forest regions in the north of France. The turning point was reached earlier in the forest regions located in the south and west of France characterised by an oceanic climate and later for the forest regions in the north and east of France characterised by a semi-continental climate (Fig. 2). In 69% (31 out of the 45) of the forest regions with hump-shaped trends, the predicted productivity rate at the end of the study period was found significantly lower than the productivity at the beginning (posterior probability larger than 0.9; Fig. S7). In two forest regions of northern France (region code B43 and C41, see Table S1) displaying a clear hump-shaped trend, the absolute growth rate at the end of the studied period was larger than at the beginning. In less than 50% (40) of the forest regions, the evidence of a hump-shaped trend was low. These 40 regions were mostly situated in the south and west of France (Fig. 1). There was strong evidence (posterior probability > 0.9) for negative linear trends in 32 of these; and for the other 8 there was no evidence of a linear trend in any direction, these regions were situated in the south-west (Grand Ouest, 4 forest regions out of 11 in total in this biogeographical region), the Alps (3/6) and in the Jura (1/2) mountain ranges. Exploring the potential climatic drivers for these variations in trend shapes revealed, in lowland regions, the presence of a threshold at a mean temperature of 11.9°C (Fig. 3). Forest regions with mean temperature above this threshold (to the exception of 5 forest regions) showed linear declines, while forest regions below this threshold showed hump-shaped trends. No such threshold in mean temperature discriminating between linear declines and hump-shaped trends could be found in mountainous regions. Water deficit tended to be higher in forest regions showing linear declines compared to regions showing hump-shaped trends both in lowlands and mountainous regions. Mountainous regions with water deficit below a threshold of 25mm showed no trend, i.e., their absolute growth rate remained constant over the considered period.

**Figure 2:**
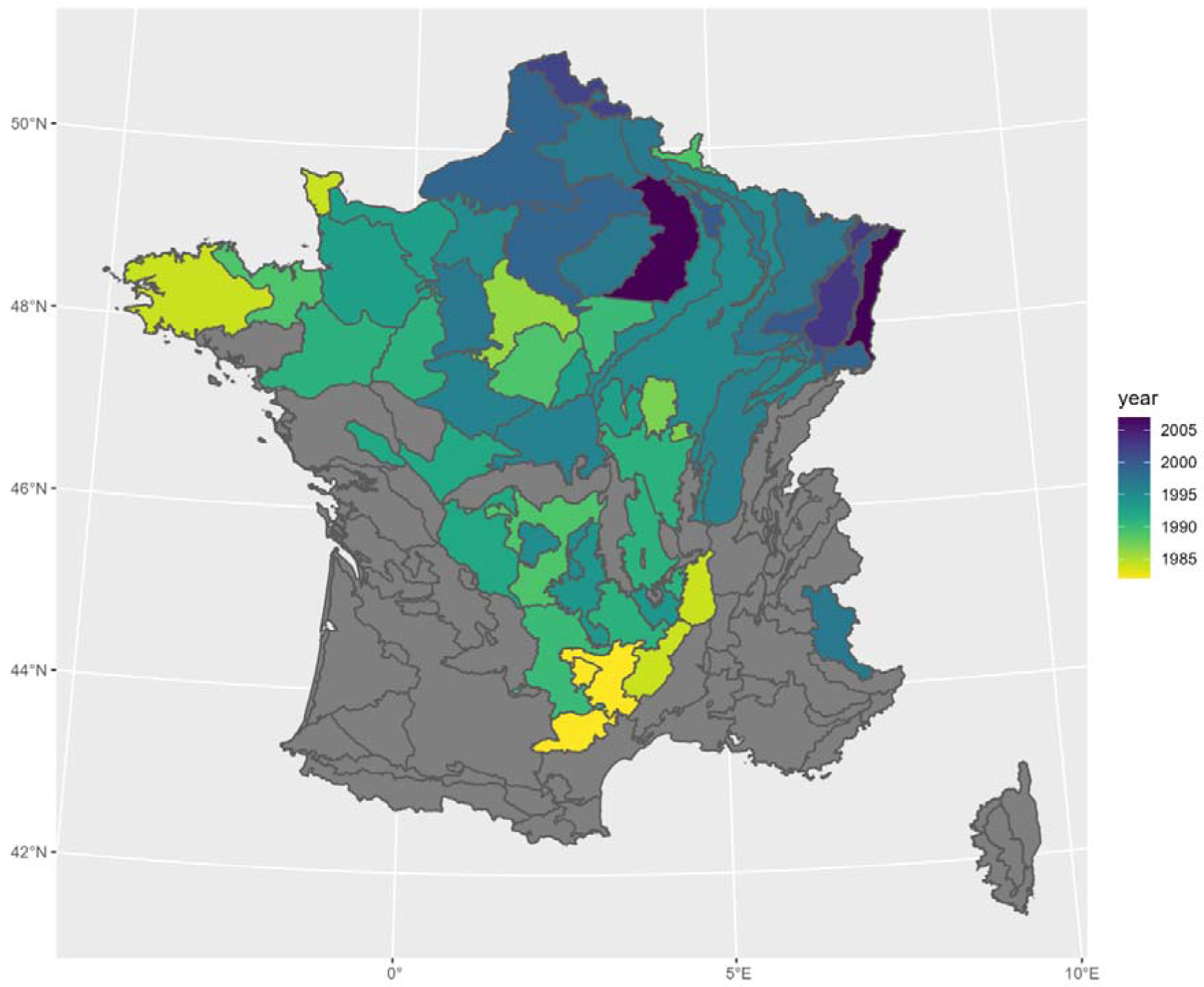
Mapping of the estimated year when the rate of productivity reached the turning point in the individual forest regions. The optimum was derived from the predicted temporal trends as shown in Fig. 1. Forest regions in grey have low evidence of having a turning point (P(β2 < 0) < 0.9).

**Figures 3:**
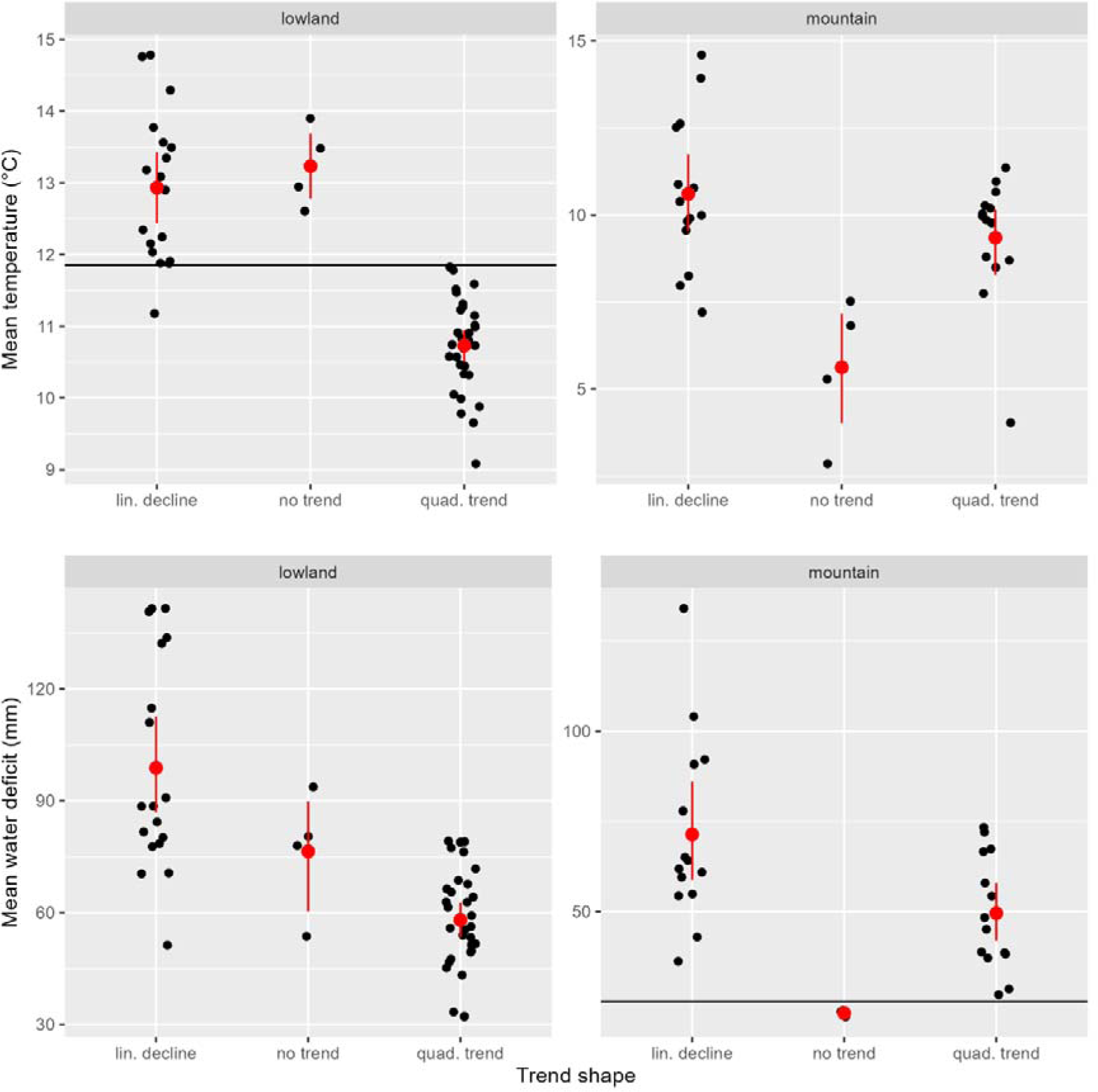
Relation between the shape of the trend in absolute growth rate at the level of the forest region and the 30-year average temperature (top row) and the water deficit (bottom row) for the reference period (1980-2009). The forest regions are separated according to their elevation in two classes: lowland and mountain. Forest regions trends are classified in three groups based on the estimated regression coefficient in the trend model: (i) no trend, P(β1 < 0) < 0.9 and P(β2 < 0) < 0.9, (ii) linear decline, P(β1 < 0) >= 0.9 and P(β2 < 0) < 0.9 and (iii) quadratic trend, P(β2 < 0) >= 0.9. Each dot is a forest region, the red dots with the vertical bars represent the mean and 95% confidence intervals.

### Climate-change effect on region-level trends

All five nested models including the first axis of a PCA on temperature anomalies and water deficit anomalies for spring and summer showed convergence for all parameters. The R² value for the full model was 0.86 when considering all model parameters and 0.30 when considering only the population-level parameters. Correlating the temporal trend to the climate models predictions mapped to the corresponding years revealed that water deficit alone poorly reproduced the temporal trends (correlation coefficient of 0.34 on average). In forest regions with linear temporal trends, temperature anomalies alone could reproduce with the highest correlation the temporal trend, more complex models including squared temperature anomalies or water deficit did not achieve higher correlation coefficients (Fig. S8). In forest regions with quadratic temporal trend, temperature anomalies plus squared temperature anomalies achieved the highest correlation to the predicted temporal trends above the model including temperature anomalies and water deficit or the full model (Fig. S8). The effect of the climate change variables on productivity is assessed within the full model. At the population-level (across all regions), there was evidence for linear negative effects of temperature (-4.87, 95% CrI -7.91, -1.76) and summer water deficit anomalies on absolute growth rates (-0.06, 95% CrI -0.11, 0.00). In other words, warmer temperature (across seasons) and drier summer months were correlated with lower absolute growth rate. The temperature anomalies linear effects were highly variable between the biogeographical and forest regions, while the water deficit had more constant effects (Fig. 4). Temperature anomalies had markedly more negative impacts on absolute growth rate in biogeographical regions situated in the south of France (Méditérranée, Pyrénnées, Corse), while biogeographical regions in the North and East of France (Centre Nord, Grand Est, Vosges) showed less negative response in absolute growth rate to increasing temperature anomalies. Two biogeographical regions in the North of France (Centre nord and Grand Est) showed evidence for hump-shaped relation with temperature anomalies. Summer water deficit anomalies tended to have more negative effect on absolute growth rate than spring water deficit anomalies with a posterior probability of 0.84. Comparing the effect sizes of the different climatic variables revealed that temperature anomalies effects were twice as large as the effect of anomalies in summer water deficit and seven times as large as the effect of anomalies in spring water deficit.

**Figure 4:**
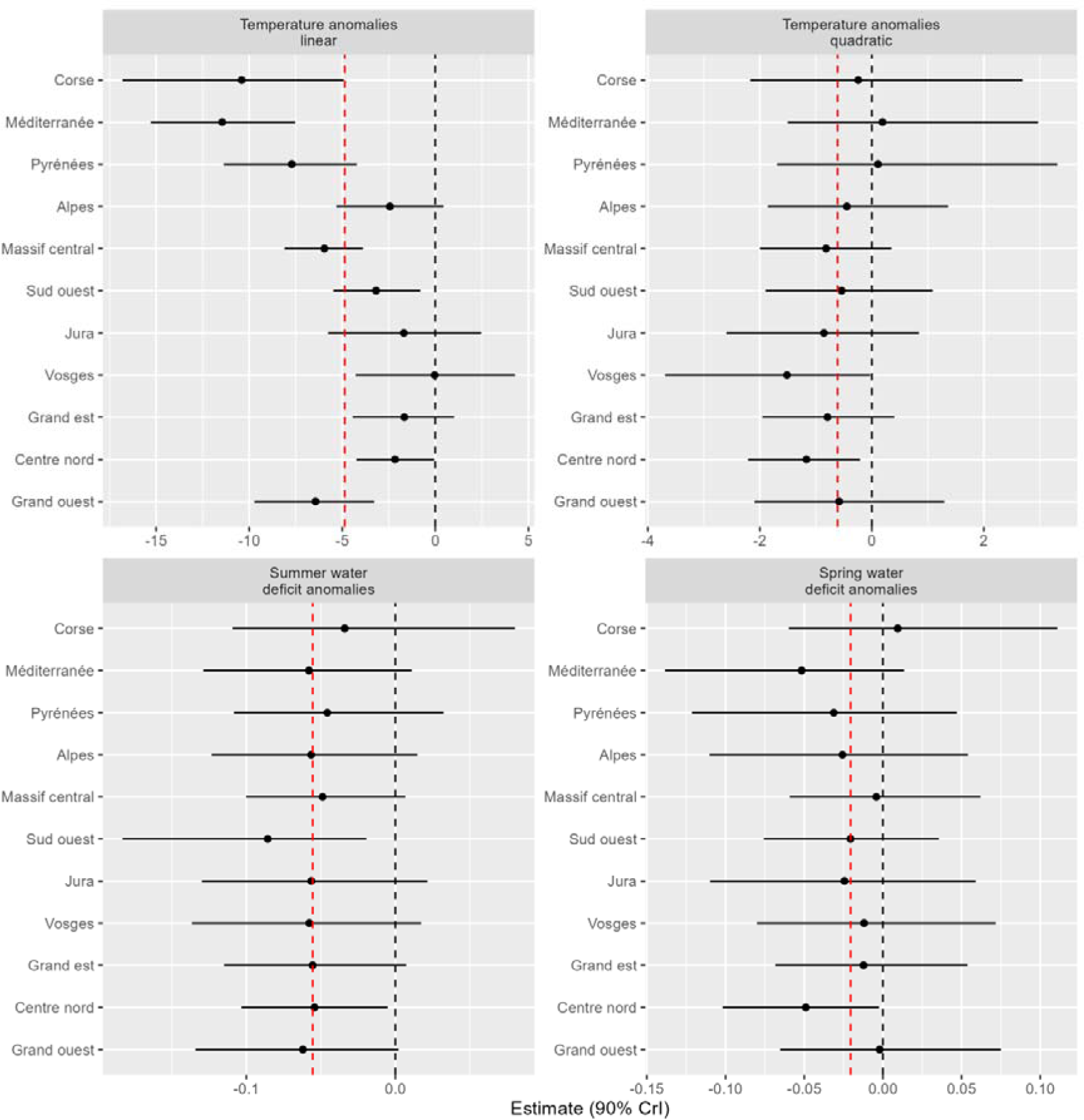
Estimated regression coefficients for the effect of the tested climatic variables at the level of the biogeographical region from the climate model defined in equation 6. The horizontal lines represent the 90% confidence intervals and the dotted vertical red line the average effect across biogeographical regions.

## Discussion

Our results showed that across climatic zones, site conditions, species composition and forest structure, French forest absolute growth rate reached a turning point and widespread declines are currently observed. These declines were found to be associated with changes in the climatic conditions mediated predominantly by the negative impact of increasing anomalies in temperature and to a lesser extent to summer water deficit.

### Variations in temporal trends between the regions

Using representative and systematically collected data on 4Mio of trees from more than 300’000 plots and more than 100 tree species our analysis revealed divergence in productivity trends depending on climate, elevation and regional contexts. Between 1980 and the early year 2000, lowland forest regions characterised by climate with average temperatures below 11.9°C, showed increases in productivity while regions situated in warmer climate, such as in the Mediterranean climatic zone, showed decline. In mountain regions, temperature and water availability did not appear to be strong determinant of trend divergence. This contrast between increasing productivity in cooler climate and declines in warmer climate has already been reported (Charru et al. 2017, Wang et al. 2023). However, this study is the first to reveal a turning point in productivity in French forests with declines throughout the country and across climatic zones after 2000. Earlier analysis on the French National Forest Inventory data (Charru et al. 2017, Ols et al. 2020) or using dendrochronological records (Spiecker et al. 1996) did not reveal widespread declines but were based on different temporal windows, on different spatial scale and were restricted to pure, even-aged stands with a limited set of species while this analysis is based on all observed forest stands. This convergence in trends in recent years is remarkable given the diverse set of tree species present in French forest, the national inventory records more than 100 different tree species. In addition, our results show that for 8 forest regions there were no evidence of a trend in any direction, these regions were situated at both ends of the normal temperature gradient (across forest regions) with warm (above 12°C) regions in the oceanic south-west of France (region code F12, F21, F23 and F51, see the Table S1 for the region names), and cold (below 7°C) and wet (water deficit below 25mm) regions in the north of the Alps (region code H10, H21 and H22) and in the Jura (region code E20) mountain regions. Forests in these mountainous regions are likely not yet negatively impacted by the recent climatic changes (Charru et al. 2017). This hypothesis is supported by the fact that none of the tested temperature and water deficit anomalies had significant impact on productivity in these regions before 2022. The four forest regions of the oceanic south-west comprise two regions with high dominance of intensive maritime pine stands (Landes), this species showed a high resistance in recent years with low rate of mortality compared to other coniferous tree species (mortality data from the French NFI). This higher resistance could be due to less marked climatic changes in these forest regions (buffering oceanic effect), or to the intrinsic capacity of this species to withstand perturbations (Rita et al. 2020, Ols et al. 2020, Ols et al. 2021). The other two forest regions are dominated by various deciduous species and show contrasted soil conditions. Reasons for the stability in productivity in these regions remain to be uncovered.

### Effect of climate change on the trends

The temporal trends could be reproduced with high fidelity in the climatic models. Investigation of nested models revealed the relative higher importance of temperature anomalies compared to water deficit to reproduce the temporal trends. The full climatic model revealed that the largest effect of a changing climate on forest productivity was the negative impact of increased temperatures, followed by the negative impact of summer water deficit. The temperature warming effect was however highly variable depending on the biogeographical context with markedly more negative effect of warmer temperature in regions of Mediterranean southern France and less pronounced, or even insignificant, effect in regions of northern France in various climatic context. In two biogeographical regions Centre Nord (northern oceanic plains) and Vosges (mid-altitude mountains), there were evidences of hump-shaped relation between productivity and temperature anomalies indicating that the impact of warming temperature on these forests where first positive but are now negative. Warming effects on forest productivity are therefore not equal across large latitudinal or climatic gradients (see also Wang et al. 2023 and Babst et al. 2019). However, as climate warming proceed in France and Europe, higher temperature and heat extremes will further impact the productive capacities of forests (EEA 2023). For instance, air temperature above 40°C that were largely absent from France in the past, will become more frequent in the future. These extreme temperatures disrupt the photosynthetic machinery of plants that have not developed adaptation mechanism to these heat stresses (Teskey et al. 2014) and lead to reduced productivity (van der Woude et al. 2023). Temperature effects are also tightly linked to water availability especially in the hotter and drier context of the Mediterranean climatic zone (Diaz-Martinez et al. 2023). The more negative impact of warming in these regions is likely to be linked to the concurrent water limitation in these contexts. Compared to the large regional variations in the effect of warming, seasonal water deficit anomalies had markedly more constant impact on forest productivity. Disruption in precipitation regimes, and in soil water availability, with on-going climate change is therefore likely to have similar effect on forest productivity across climate zone (but see Rita et al. 2020). Our results confirmed the negative impact of summer and spring water deficit on forest productivity across forest regions reported in previous studies (Ols et al. 2020). We also found that anomalies in summer water deficit had stronger effect than spring ones, outlining the stronger water-limitation of productivity especially in summer month in these forests. More researches and meta-analysis should evaluate if disruption in soil moisture regimes with on-going climate change leads to similar effect on forest productivity across climate zone.

### Caveats

The results presented here suffer from various limitations and caveats that are outlined below. First, because the French NFI has relied solely on temporary plots until 2010, our data only takes into account the stocks and productivity of living trees, ignoring harvested and dead trees. Changes in the methodology of the French NFI allows to take into account dead and harvested trees starting from the year 2010 and future analysis could derive trends in gross productivity considering all trees. We could show for the years 2010 until 2017 that the estimations of living trees productivity were tightly correlated to the estimations considering all trees. However, future increase in stress and further pest outbreaks could lead to other patterns with mortality and harvest also touching the most vigorous individuals. Second, the estimation of productivity is based on tree cores directly measured in the fields (IGN, 2023), that are prone to measurement errors in certain species (i.e. evergreen oak). The protocol to estimate productivity being constant through the considered temporal window, we assume that these errors are temporally constant and are not biasing the estimated trends and associations with climate change. Another caveat is our use of a coarse variable to quantify and model the impact of changes in temperature. Given the high level of correlations between the 12 derived temperature variables, we chose to synthetise the information by using only one PCA axis. This approach prevents us to pinpoint the major seasonal and distributional (mean vs. minimum vs. maximum) determinant linking temperature changes and forest productivity.

## Conclusion

Following the reported increase in forest productivity over the 20^th^ century, our results show that a turning point has been reached and widespread declines are currently observed in 95% of the regions with variable tree species composition and forest structure. This result highlights the limited potential of actively managing tree species composition with already present phenotypes and species to boost forest productivity within the current forestry regimes. Our analysis at the regional level revealed variations in trend shapes mediated by mean temperature and water availability in lowland regions, while variation in trend shapes in mountain regions appears to be driven by other factors. The derived temporal trends correlated with climate change throughout France which has several implications for sylvicultural management but also for public policies. Declining productivity implies that extraction rate of timber should decline and rotation time in regular stands should increase to ensure the sustainability of management practices. Public policy at the EU and French level strive to bring net CO_2_ emission to 0 by 2050, these policies rely on the capacity of natural ecosystems, including forests, to capture and store CO_2_ for long period of times. Declining productivity will lead to declining carbon sinks if ecosystem respiration and decomposition rates do not mirror the productivity declines (Ciais et al. 2005) and if forests are simultaneously affected by disturbances such as fires or storms leading to pervasive shifts in forest ecosystems (Mc Dowell et al. 2020). National accounting of greenhouse gases reported already the declining carbon sinks of forests (EEA 2023). These policies should therefore also consider the uncertainties around the capacity of forests and natural ecosystem to store CO_2_ under climatic change.

## Data availability statement

Data and code to reproduce the models and the figures are archived under https://zenodo.org/doi/10.5281/zenodo.10836876

