## Supplementary material for "Turning point in forest productivity revealed from 40 years of national forest inventory data": SI

List of the supporting information materials:

- Table S1: List of 86 forest regions with their respective names, biogeographical region and dominant tree species
- Fig. S1: Scatterplot of the estimated rate of productivity considering all trees (gross) against rate of productivity considering only living trees (net).
- Table S2: Climatic variables loadings on the first three PC axis
- Table S3: Priors used on model parameters
- Fig. S2: Trend in growing stock
- Fig. S3: Trend in productivity
- Fig. S4: Trend in quadratic mean diameter
- Fig. S5: Residual vs fitted values from the quadratic model per regions
- Fig S6: Residuals vs covariates and residual posterior distribution from the quadratic model
- Table S4: Variation in model coefficients between biogeographical regions and forest regions
- Fig. S7: Mapping of the predicted difference between productivity at the end and at the beginning of the temporal window (1978-2022)
- Fig. S8: Correlation between the temporal trend predicted from the model defined in equations 1-4 and predictions from the nested climate models
- Fig. S9: Correlation between the trend and the climatic model.

Table S1: List of 86 forest regions with their respective names, biogeographical region and dominant tree species

| Code | Name | Biog. Region | Dominant species |
| --- | --- | --- | --- |
| A11 | Ouest-Bretagne et Nord-Cotentin | Grand ouest | Quercus robur/Picea sitchensis/Fagus sylvatica/Castanea sativa/Pseudotsuga menziesii/Betula pubescens/Quercus petraea/Pinus pinaster |
| A12 | Pays de Saint-Malo | Grand ouest | Quercus robur/Castanea sativa/Pinus pinaster/Quercus petraea/Pinus sylvestris/Fagus sylvatica/Picea sitchensis/Populus ident |
| A13 | Bocage normand et Pays de Fougères | Grand ouest | Quercus petraea/Fagus sylvatica/Quercus robur/Pseudotsuga menziesii/Castanea sativa/Pinus sylvestris/Betula pubescens |
| A21 | Bretagne méridionale | Grand ouest | Pinus pinaster/Quercus robur/Castanea sativa/Fagus sylvatica/Betula pubescens/Pinus sylvestris |
| A22 | Bocage armoricain | Grand ouest | Quercus petraea/Quercus robur/Castanea sativa/Pinus pinaster/Pinus sylvestris/Fagus sylvatica/Pseudotsuga menziesii |
| A30 | Bocage vendéen | Grand ouest | Quercus robur/Quercus petraea/Populus ident/Castanea sativa/Pinus pinaster/Fraxinus excelsior/Pinus ident |
| B10 | Côtes et plateaux de la Manche | Centre nord | Fagus sylvatica/Quercus robur/Quercus petraea/Fraxinus excelsior/Carpinus betulus/Acer pseudoplatanus/Populus ident/Betula pubescens/Castanea sativa |
| B21 | Flandres | Centre nord | Quercus robur/Populus ident/Fraxinus excelsior/Carpinus betulus/Salix ident/Betula pubescens/Pinus sylvestris/Quercus petraea |
| B22 | Plaine picarde | Centre nord | Populus ident/Fraxinus excelsior/Acer pseudoplatanus/Quercus robur/Fagus sylvatica/Carpinus betulus/Tilia cordata/Salix ident/Alnus glutinosa/Betula pubescens |
| B23 | Mosan, Thiérache et Hainaut | Centre nord | Quercus robur/Carpinus betulus/Fraxinus excelsior/Quercus petraea/Fagus sylvatica/Acer pseudoplatanus/Populus ident |
| B31 | Campagne de Caen et Pays d'Auge | Centre nord | Quercus petraea/Quercus robur/Fagus sylvatica/Fraxinus excelsior/Pseudotsuga menziesii/Betula pubescens/Populus ident/Alnus glutinosa/Populus tremula/Castanea sativa |
| B32 | Plateaux de l'Eure | Centre nord | Quercus petraea/Quercus robur/Fagus sylvatica/Carpinus betulus/Pseudotsuga menziesii/Pinus sylvestris/Betula pubescens |
| B33 | Perche | Centre nord | Quercus petraea/Quercus robur/Castanea sativa/Carpinus betulus/Fagus sylvatica |
| B41 | Bassin parisien tertiaire | Centre nord | Quercus petraea/Quercus robur/Fagus sylvatica/Castanea sativa/Fraxinus excelsior/Carpinus betulus/Pinus sylvestris/Populus ident |
| B42 | Brie et Tardenois | Centre nord | Quercus robur/Quercus petraea/Fraxinus excelsior/Carpinus betulus/Populus tremula/Betula pubescens |
| B43 | Champagne crayeuse | Centre nord | Fraxinus excelsior/Populus ident/Pinus sylvestris/Pinus nigra/Quercus robur/Acer pseudoplatanus/Alnus glutinosa/Quercus petraea/Betula pubescens/Salix ident Quercus petraea/Quercus robur/Fraxinus |
| B44 | Beauce | Centre nord | excelsior/Carpinus betulus/Quercus pubescens/Robinia pseudoacacia/Pinus sylvestris/Populus ident/Castanea sativa/Pinus nigra |
| B51 | Champagne humide | Centre nord | Quercus robur/Quercus petraea/Carpinus betulus/Fraxinus excelsior/Populus tremula/Populus ident |
| B52 | Pays d'Othe et Gatinais oriental | Centre nord | Quercus petraea/Quercus robur/Carpinus betulus/Robinia pseudoacacia/Pseudotsuga menziesii |
| B53 | Pays-Fort, Nivernais et plaines prémorvandelles | Centre nord | Quercus petraea/Quercus robur/Carpinus betulus/Fagus sylvatica |
| B61 | Baugeois-Maine | Centre nord | Pinus pinaster/Castanea sativa/Quercus robur/Quercus petraea/Populus ident |
| B62 | Champeigne-Gâtine tourangelle | Centre nord | Quercus petraea/Quercus robur/Pinus pinaster/Castanea sativa/Carpinus betulus/Populus ident |
| B70 | Sologne-Orléanais | Centre nord | Quercus robur/Pinus sylvestris/Quercus petraea/Pinus ident/Betula pubescens |
| B81 | Loudunais et Saumurois | Centre nord | Quercus robur/Pinus pinaster/Quercus pubescens/Populus ident/Quercus petraea/Castanea sativa |
| B82 | Brenne et Brandes | Centre nord | Quercus robur/Quercus petraea/Quercus pubescens/Pinus pinaster/Carpinus betulus |
| B91 | Boischaut et Champagne berrichonne | Centre nord | Quercus petraea/Quercus robur/Carpinus betulus |
| B92 | Bourbonnais et Charolais | Centre nord | Quercus petraea/Quercus robur/Pseudotsuga menziesii |
| C11 | Ardenne primaire | Grand est | Picea abies/Quercus petraea/Betula pubescens/Quercus robur |
| C12 | Argonne | Grand est | Quercus petraea/Picea abies/Fagus sylvatica/Carpinus betulus/Quercus robur/Pseudotsuga menziesii |
| C20 | Plateaux calcaires du Nord-Est | Grand est | Quercus petraea/Fagus sylvatica/Carpinus betulus/Quercus robur/Fraxinus excelsior/Picea abies/Pinus sylvestris/Acer campestre |
| C30 | Plaines et dépressions argileuses du Nord-Est | Grand est | Quercus petraea/Quercus robur/Fagus sylvatica/Carpinus betulus/Fraxinus excelsior |
| C41 | Plaine d'Alsace | Grand est | Quercus robur/Fraxinus excelsior/Pinus sylvestris/Carpinus betulus/Quercus petraea/Alnus glutinosa/Fagus sylvatica/Robinia pseudoacacia |
| C42 | Sundgau alsacien et belfortain | Grand est | Fagus sylvatica/Fraxinus excelsior/Quercus robur/Quercus petraea/Carpinus betulus |
| C51 | Saône, Bresse et Dombes | Grand est | Quercus petraea/Quercus robur/Carpinus betulus/Fagus sylvatica/Fraxinus excelsior/Robinia pseudoacacia/Populus tremula |
| C52 | Plaines et piémonts alpins | Grand est | Castanea sativa/Fraxinus excelsior/Quercus petraea/Fagus sylvatica/Picea abies/Robinia pseudoacacia/Quercus robur/Carpinus betulus/Alnus glutinosa/Quercus pubescens/Tilia cordata |
| D11 | Massif vosgien central | Vosges | Abies alba/Picea abies/Fagus sylvatica/Pinus sylvestris |
| D12 | Collines périvosgiennes et Warndt | Vosges | Fagus sylvatica/Quercus petraea/Abies alba/Picea abies/Quercus robur/Pinus sylvestris |
| E10 | Premier plateau du Jura | Jura | Fagus sylvatica/Abies alba/Picea abies/Quercus petraea/Fraxinus excelsior/Carpinus betulus/Quercus robur |
| E20 | Deuxième plateau et Haut-Jura | Jura | Picea abies/Abies alba |
| F11 | Terres rouges | Sud ouest | Castanea sativa/Quercus robur/Quercus petraea/Carpinus betulus/Quercus pubescens |
| F12 | Groies | Sud ouest | Quercus pubescens/Acer campestre/Quercus robur/Fraxinus excelsior/Populus ident/Quercus petraea |
| F13 | Marais littoraux | Sud ouest | Populus ident |
| F14 | Champagne charentaise | Sud ouest | Quercus robur/Quercus pubescens/Castanea sativa/Populus ident/Fraxinus excelsior/Pinus pinaster/Carpinus betulus |
| F15 | Périgord | Sud ouest | Castanea sativa/Quercus pubescens/Quercus robur/Pinus pinaster/Carp |
| F21 | Landes de Gascogne | Sud ouest | Pinus pinaster |
| F22 | Dunes atlantiques | Sud ouest | Pinus pinaster |
| F23 | Bazadais, Double et Landais | Sud ouest | Pinus pinaster/Quercus robur/Castanea sativa |
| F30 | Coteaux de la Garonne | Sud ouest | Quercus pubescens/Quercus robur/Quercus petraea/Castanea sativa/Carpinus betulus/Pinus pinaster/Fraxinus excelsior/Populus ident/Robinia pseudoacacia |
| F40 | Causses du Sud-Ouest | Sud ouest | Quercus pubescens/Acer campestre |
| F51 | Adour atlantique | Sud ouest | Quercus robur/Alnus glutinosa/Castanea sativa/Robinia pseudoacacia/Fraxinus excelsior/Pinus pinaster |
| F52 | Collines de l'Adour | Sud ouest | Quercus robur/Castanea sativa/Alnus glutinosa/Fraxinus excelsior/Pinus pinaster/Populus ident/Robinia pseudoacacia |
| G11 | Châtaigneraie du Centre et de l'Ouest | Massif central | Quercus robur/Castanea sativa/Pseudotsuga menziesii/Fagus sylvatica/Carpinus betulus/Betula pubescens/Quercus petraea |
| G12 | Marches du Massif central | Massif central | Quercus robur/Quercus petraea/Pseudotsuga menziesii/Carpinus betulus/Castanea sativa |
| G13 | Plateaux limousins | Massif central | Quercus robur/Fagus sylvatica/Pseudotsuga menziesii/Castanea sativa/Picea abies/Pinus sylvestris/Quercus petraea |
| G21 | Plateaux granitiques ouest du Massif central | Massif central | Pseudotsuga menziesii/Picea abies/Fagus sylvatica/Quercus robur/Abies alba |
| G22 | Plateaux granitiques du centre du Massif central | Massif central | Abies alba/Pinus sylvestris/Picea abies/Pseudotsuga menziesii |
| G23 | Morvan et Autunois | Massif central | Pseudotsuga menziesii/Quercus petraea/Picea abies/Fagus sylvatica/Quercus robur |
| G30 | Massif central volcanique | Massif central | Fagus sylvatica/Picea abies/Abies alba/Pinus sylvestris |
| G41 | Bordure Nord-Est du Massif central | Massif central | Pseudotsuga menziesii/Abies alba/Quercus petraea/Castanea sativa |
| G42 | Monts du Vivarais et du Pilat | Massif central | Abies alba/Pseudotsuga menziesii/Pinus sylvestris/Castanea sativa/Picea abies/Quercus pubescens |
| G50 | Ségala et Châtaigneraie auvergnate | Massif central | Castanea sativa/Quercus robur/Quercus petraea/Fagus sylvatica/Pseudotsuga menziesii/Quercus pubescens |
| G60 | Grands Causses | Massif central | Pinus sylvestris/Pinus nigra/Quercus pubescens |
| G70 | Cévennes | Massif central | Castanea sativa/Pinus pinaster/Fagus sylvatica/Pinus ident/Quercus ilex/Pseudotsuga menziesii/Pinus sylvestris |
| G80 | Haut-Languedoc et Lévézou | Massif central | Fagus sylvatica/Pseudotsuga menziesii/Picea abies/Castanea sativa/Abies alba/Quercus petraea/Pinus ident/Quercus robur |
| G90 | Plaines alluviales et piémonts du Massif central | Massif central | Quercus petraea/Pinus sylvestris/Quercus robur/Pseudotsuga menziesii/Fraxinus excelsior/Quercus pubescens/Abies alba/Fagus sylvatica |
| H10 | Préalpes du Nord | Alpes | Picea abies/Fagus sylvatica |
| H21 | Alpes externes du Nord | Alpes | Picea abies/Abies alba/Fagus sylvatica/Fraxinus excelsior |
| H22 | Alpes internes du Nord | Alpes | Picea abies/Larix decidua/Abies alba/Fagus sylvatica/Fraxinus excelsior |
| H30 | Alpes externes du Sud | Alpes | Pinus sylvestris/Pinus nigra/Fagus sylvatica |
| H41 | Alpes intermédiaires du Sud | Alpes | Pinus sylvestris/Larix decidua/Abies alba/Fagus sylvatica/Picea abies |
| H42 | Alpes internes du Sud | Alpes | Larix decidua/Pinus sylvestris/Abies alba |
| I11 | Marches pyrénéennes | Pyrénnés | Quercus robur/Castanea sativa/Fagus sylvatica/Quercus pubescens/Fraxinus excelsior/Quercus petraea/Robinia pseudoacacia/Pseudotsuga menziesii |
| I12 | Pyrénées cathares | Pyrénnés | Abies alba/Fagus sylvatica/Pinus sylvestris/Quercus pubescens |
| I13 | Corbières | Pyrénnés | Pinus nigra/Quercus pubescens/Quercus ilex/Fagus sylvatica/Pseudotsuga menziesii/Cedrus atlantica/Castanea sativa |
| I21 | Haute-chaîne pyrénéenne | Pyrénnés | Fagus sylvatica/Abies alba/Fraxinus excelsior/Castanea sativa/Quercus robur |
| I22 | Pyrénées catalanes | Pyrénnés | Pinus uncinata/Pinus sylvestris/Fagus sylvatica/Abies alba/Quercus pubescens/Castanea sativa |
| J10 | Garrigues | Méditerranée | Quercus ilex/Quercus pubescens/Pinus halepensis/Populus alba |
| J21 | Roussillon | Méditerranée | Quercus ilex/Castanea sativa/Quercus suber/Quercus pubescens |
| J22 | Plaines et collines rhodaniennes et languedociennes | Méditerranée | Quercus pubescens/Pinus halepensis/Quercus ilex/Populus alba/Pinus sylvestris/Fagus sylvatica/Pinus nigra |
| J23 | Provence calcaire | Méditerranée | Pinus halepensis/Quercus pubescens |
| J24 | Secteurs niçois et préligure | Méditerranée | Pinus halepensis/Quercus ilex/Quercus pubescens/Pinus pinaster |
| J30 | Maures et Esterel | Méditerranée | Quercus suber/Pinus pinaster/Quercus pubescens/Quercus ilex |
| J40 | Préalpes du Sud | Méditerranée | Quercus pubescens/Pinus halepensis/Pinus sylvestris/Quercus ilex |
| K11 | Corse occidentale | Corse | Quercus ilex/Pinus pinaster |
| K12 | Montagne corse | Corse | Pinus ident/Pinus pinaster/Quercus ilex |
| K13 | Corse orientale | Corse | Quercus ilex/Castanea sativa/Pinus pinaster/Fagus sylvatica/Arbutus unedo |

Figure S1: Scatterplot of the estimated rate of productivity considering all trees (gross) against rate of productivity considering only living trees (net). Each dot represents the estimated value for one forest region in a given temporal window. The line represents the 1:1 relation.

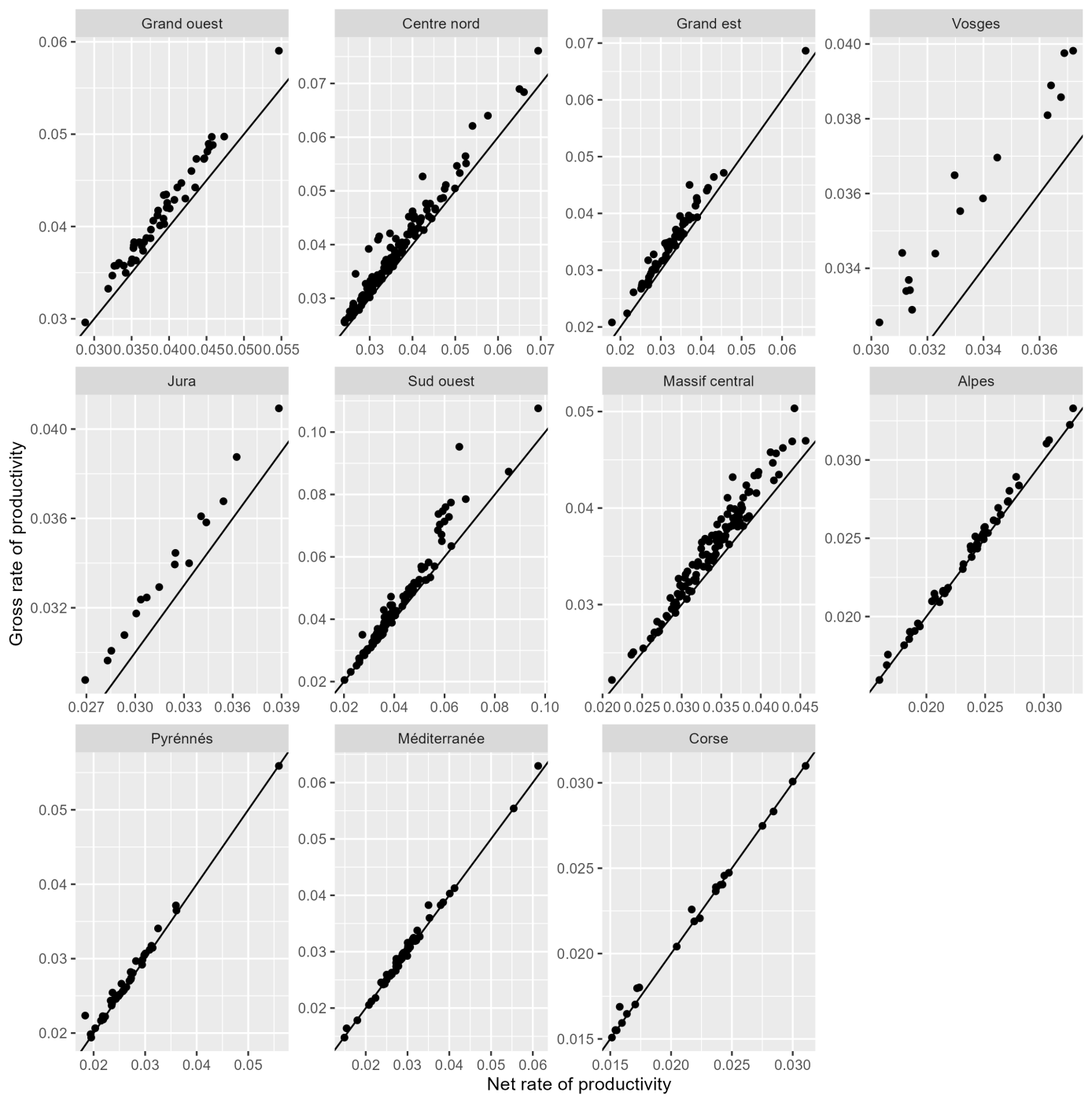

Method:

We computed for these years the gross rate of productivity computed as:

$$GR= \frac{PL+PH}{VL+VD+VH}$$

Where PL is the productivity of living trees, PH the productivity of harvested trees, VL the volume of living trees, VD the volume of dead trees and VH the volume of harvested trees. Dead trees are assumed to have low productivity in the 5 year before their death. Gross and net (PL / VL) rate of productivity were tightly correlated (Fig. S1), we will therefore use only the net rate of productivity thereafter (only considering living trees).

Table S2: PCA loadings of the temperature anomalies on the first principal axis explaining 73% of the total variation

| variables | loading |
| --- | --- |
| tminautumn | 0.26 |
| tminspring | 0.29 |
| tminsummer | 0.29 |
| tminwinter | 0.29 |
| tmaxautumn | 0.24 |
| tmaxspring | 0.32 |
| tmaxsummer | 0.3 |
| tmaxwinter | 0.3 |
| tmoyautumn | 0.26 |
| tmoyspring | 0.31 |
| tmoysummer | 0.3 |
| tmoywinter | 0.3 |

Table S3: Priors used in all fitted models, the parameter names correspond to the equations 1-4

| Parameter | Prior |
| --- | --- |
| β_0*_ | Student(3, 3.4, 2.5) |
| β_1*_ | flat |
| γ | flat |
| σ | Student(3, 0, 2.5) lower bounded at 0 |

Figure S2: Trends in volume of living trees volume estimated per forest regions and per 5-year temporal window, one dot represents a forest region for a given 5-year temporal window. A loess smooth is added per biogeographical regions and the dashed red line represent the overall trend. Note the varying y-axis range between the biogeographical region.

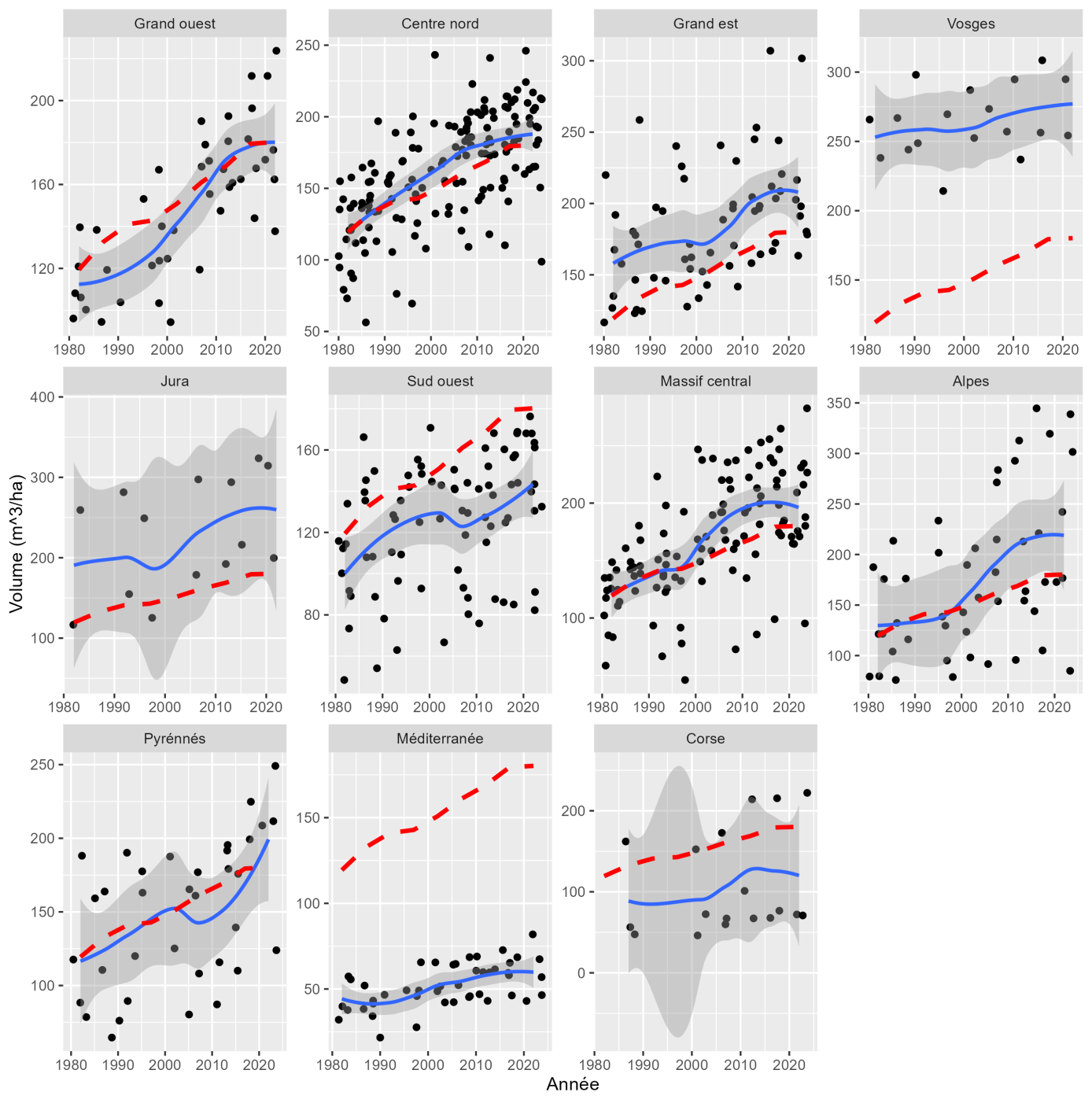

Figure S3: Trends in production of volume of living trees volume estimated per forest regions and per 5-year temporal window, one dot represents a forest region for a given 5-year temporal window. A loess smooth is added per biogeographical regions and the dashed red line represent the overall trend. Note the varying y-axis range between the biogeographical region.

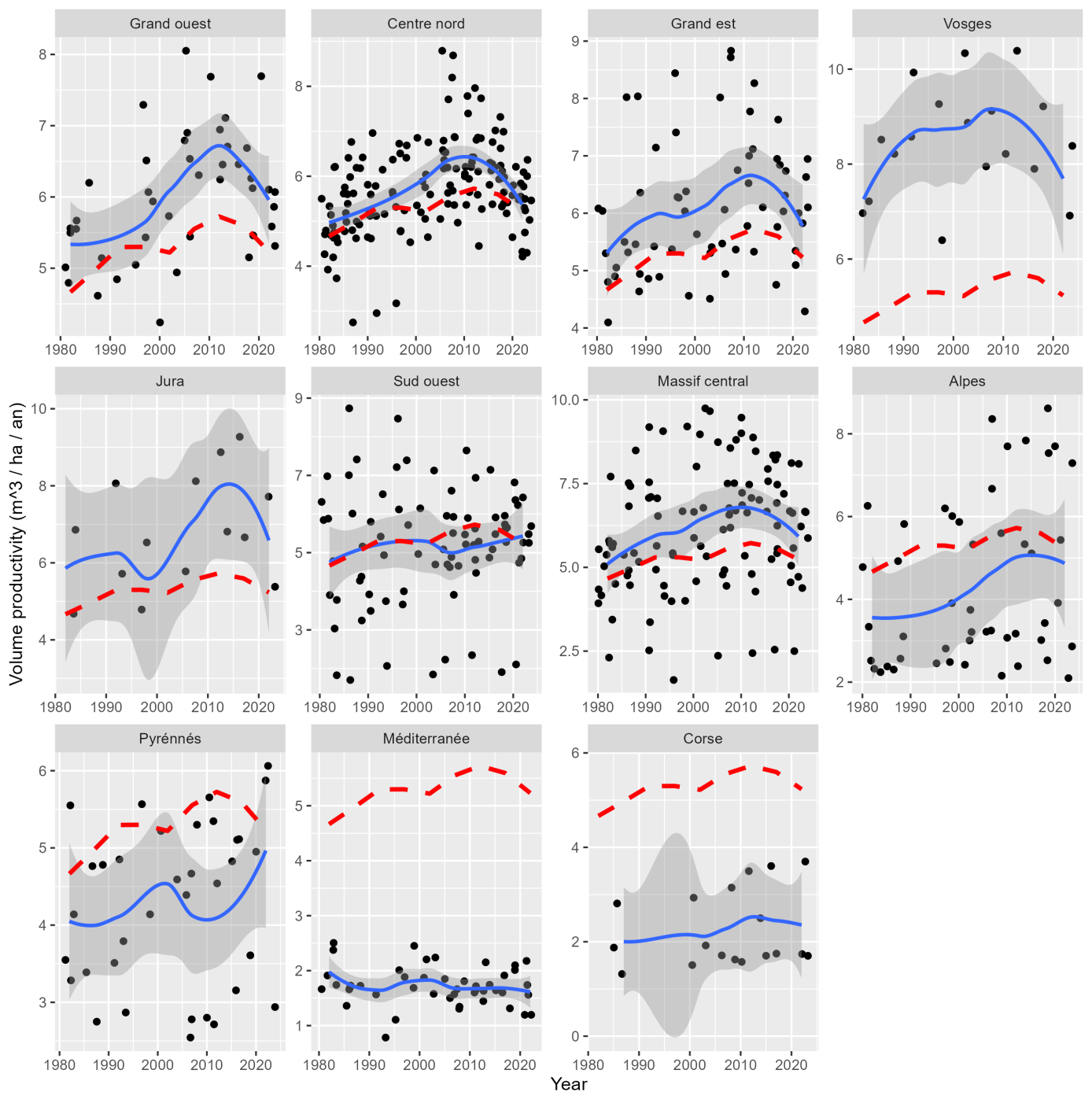

Figure S4: Trends in quadratic mean diameter (proxy for forest development stage) estimated per forest regions and per 5-year temporal window, one dot represents a forest region for a given 5-year temporal window. A loess smooth is added per biogeographical regions and the dashed red line represent the overall trend. Note the varying y-axis range between the biogeographical region.

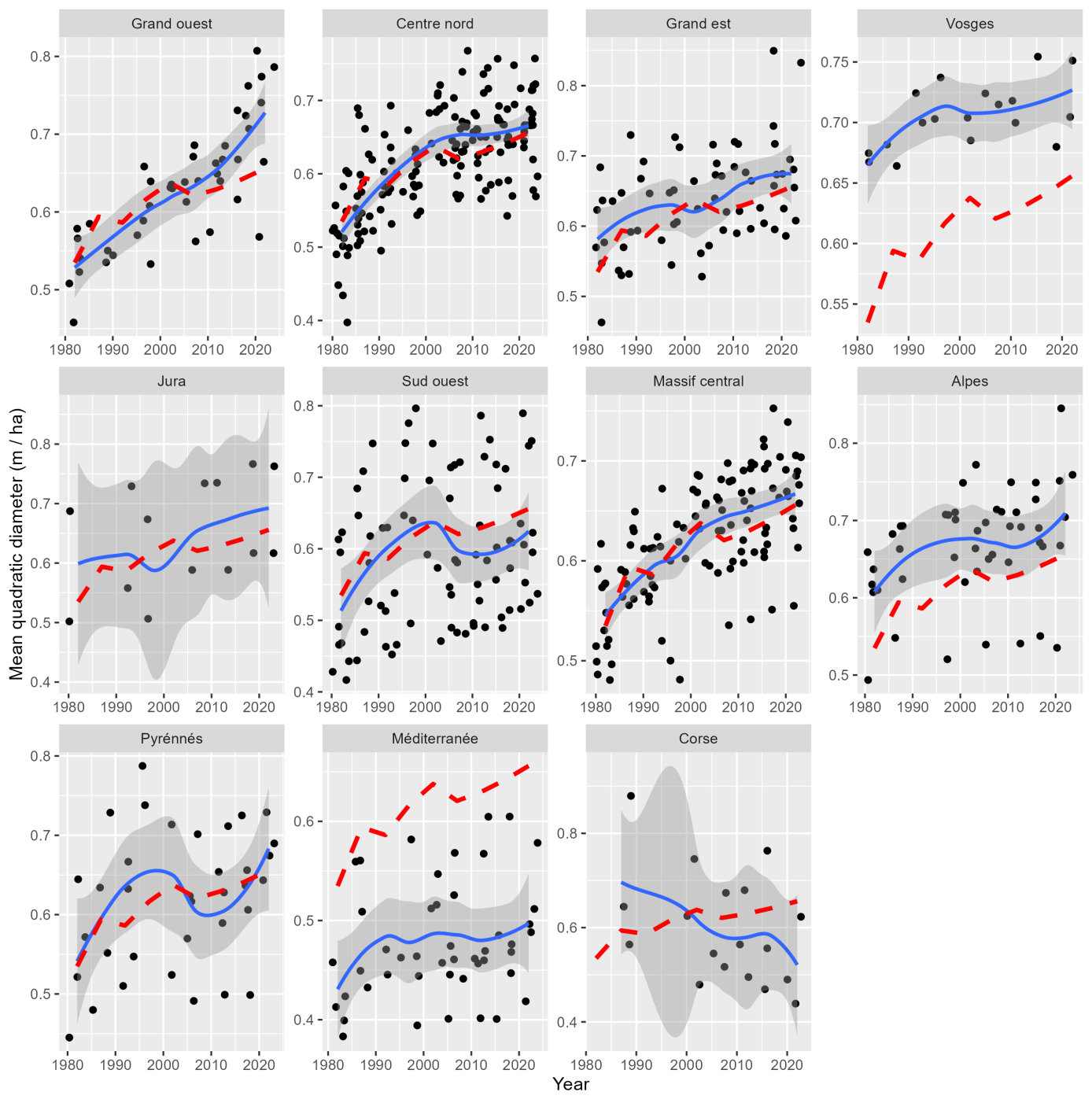

Figure S5: Residuals against the fitted values from the model defined in equations 1-4 per biogeographical regions. The vertical and horizontal bars represent the 90% Credible intervals and the horizontal dotted line is at 0.

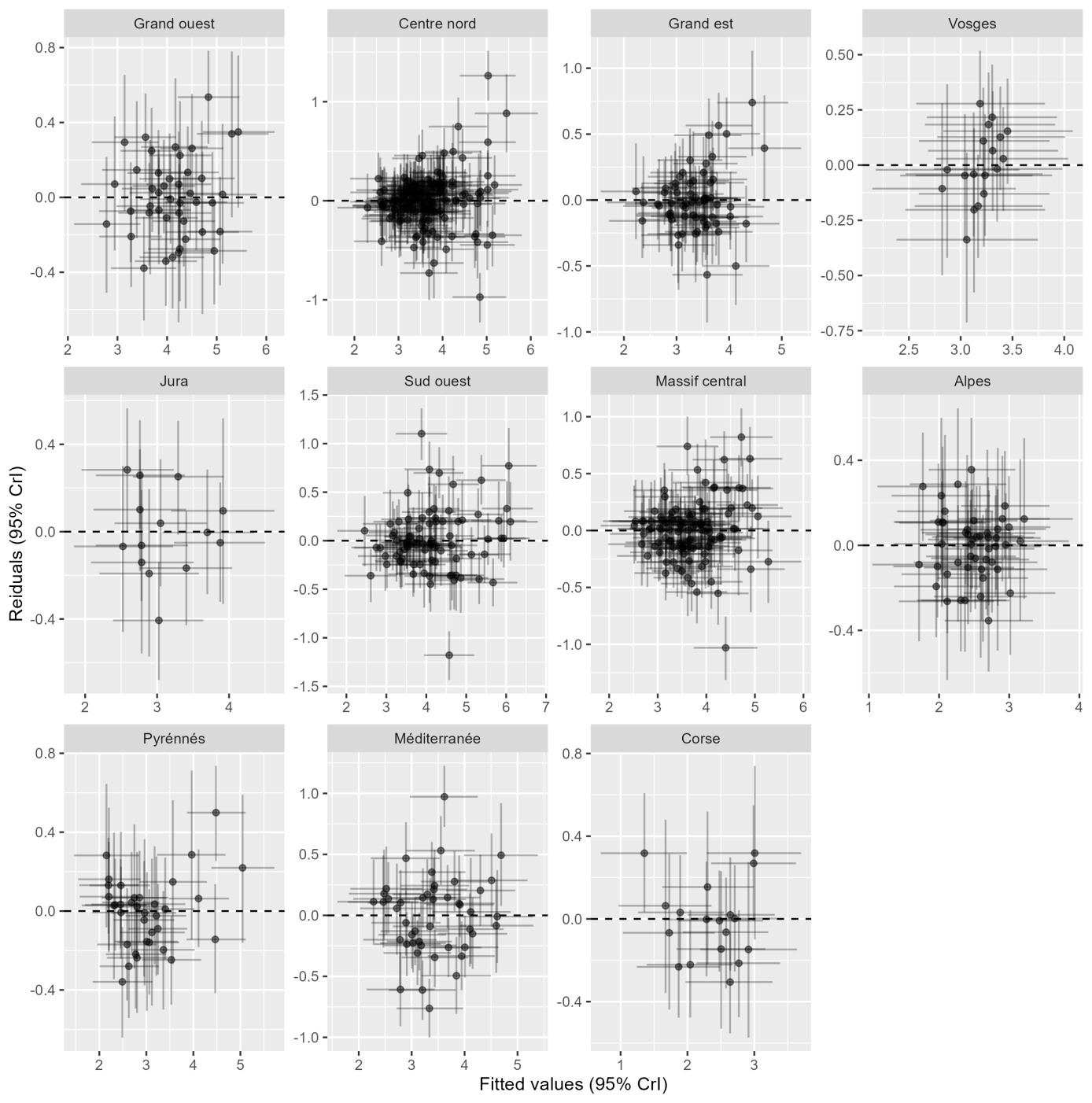

Figure S6: Plots of the residuals of the model described in equations 1-4 against the covariates in A-B, against the wood volume in C and posterior predicted distribution against the observed distribution in D.

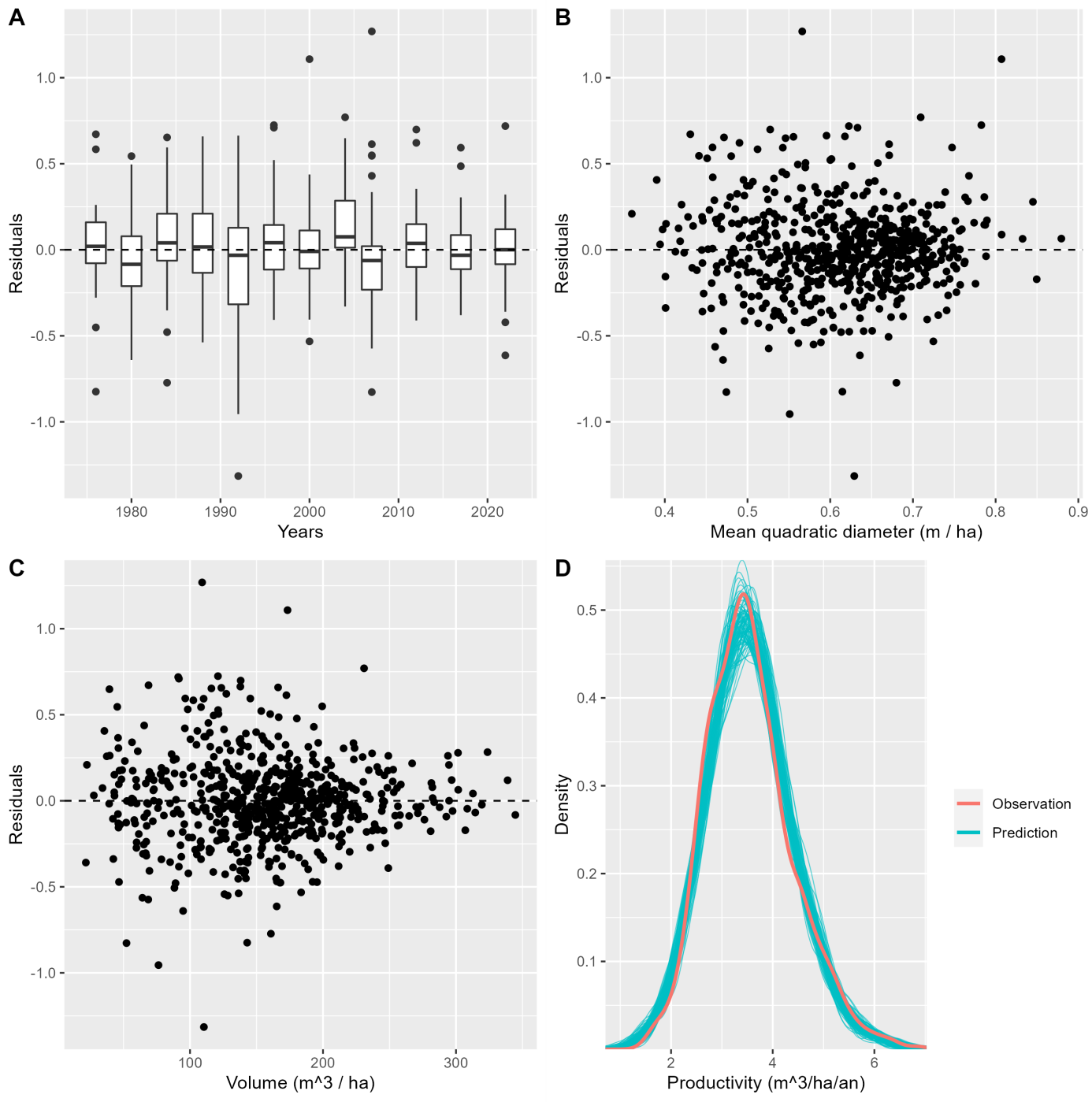

Table S4: Percent of deviance for the different hierarchical terms at the level of the regions (middle) and the level of the forest regions within the regions (right). The values are the ratio between the standard deviation of the coefficient at the given level and the sum of all standard deviation terms (including residual standard deviation). Larger values indicate larger variations in the given coefficients at the given level.

| Coefficients | Biogeographical regions (%) | Forest regions within biogeographical regions (%) |
| --- | --- | --- |
| Intercept | 1 | 0.7 |
| Linear trend | 40 | 38 |
| Quadratic trend | 9 | 9 |

Fig. S7: Predicted difference in productivity at the level of the forest regions between the end (2022) and the beginning of the temporal window (1978). Negative values indicate decline in productivity over the studied period. Regions in grey showed no significant (at the 90% level) differences in productivity.

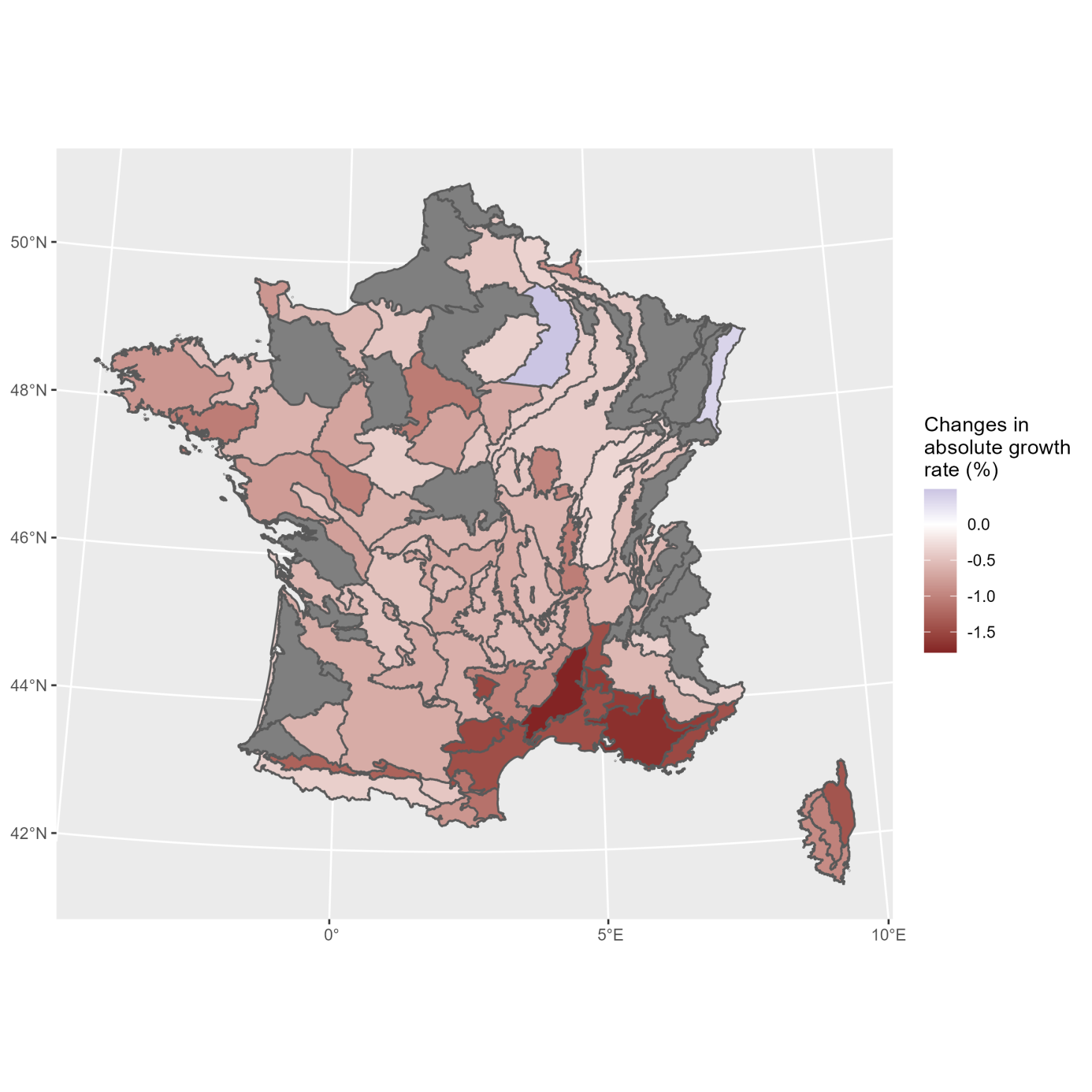

Figure S8: Correlation between the temporal trend predicted from the model defined in equations 1-4 and predictions from the nested climate models mapped to the corresponding years. Each dot represents one forest region. Forest regions are separated according to the probability of quadratic trends (p >= 0.9). The x axis represents the different nested models with (from left to right): (i) only water deficit (CWD, both summer and spring), (ii) only temperature anomalies (T), (iii) temperature anomalies and water deficit (T+CWD), (iv) temperature anomalies with quadratic term (T+T²), (iv) full model (T+CWD+T²). The diamonds represent the average correlation across forest regions and the vertical bars the 95% confidence intervals.

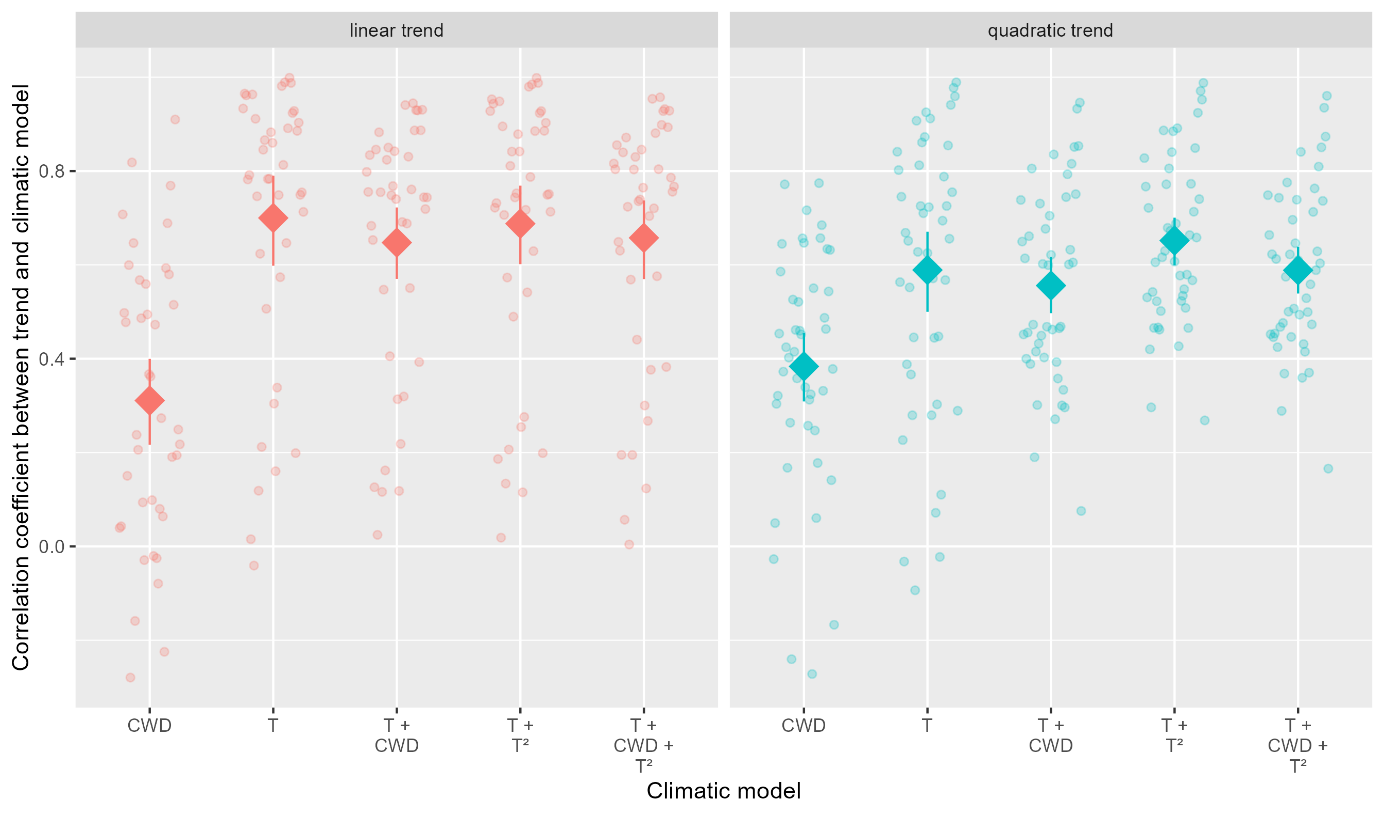

Figure S9: Scatterplot of the prediction from the model defined in equations 1-4 against the prediction from the model defined in equations 6, the vertical and horizontal bars represent the 90% credible intervals.

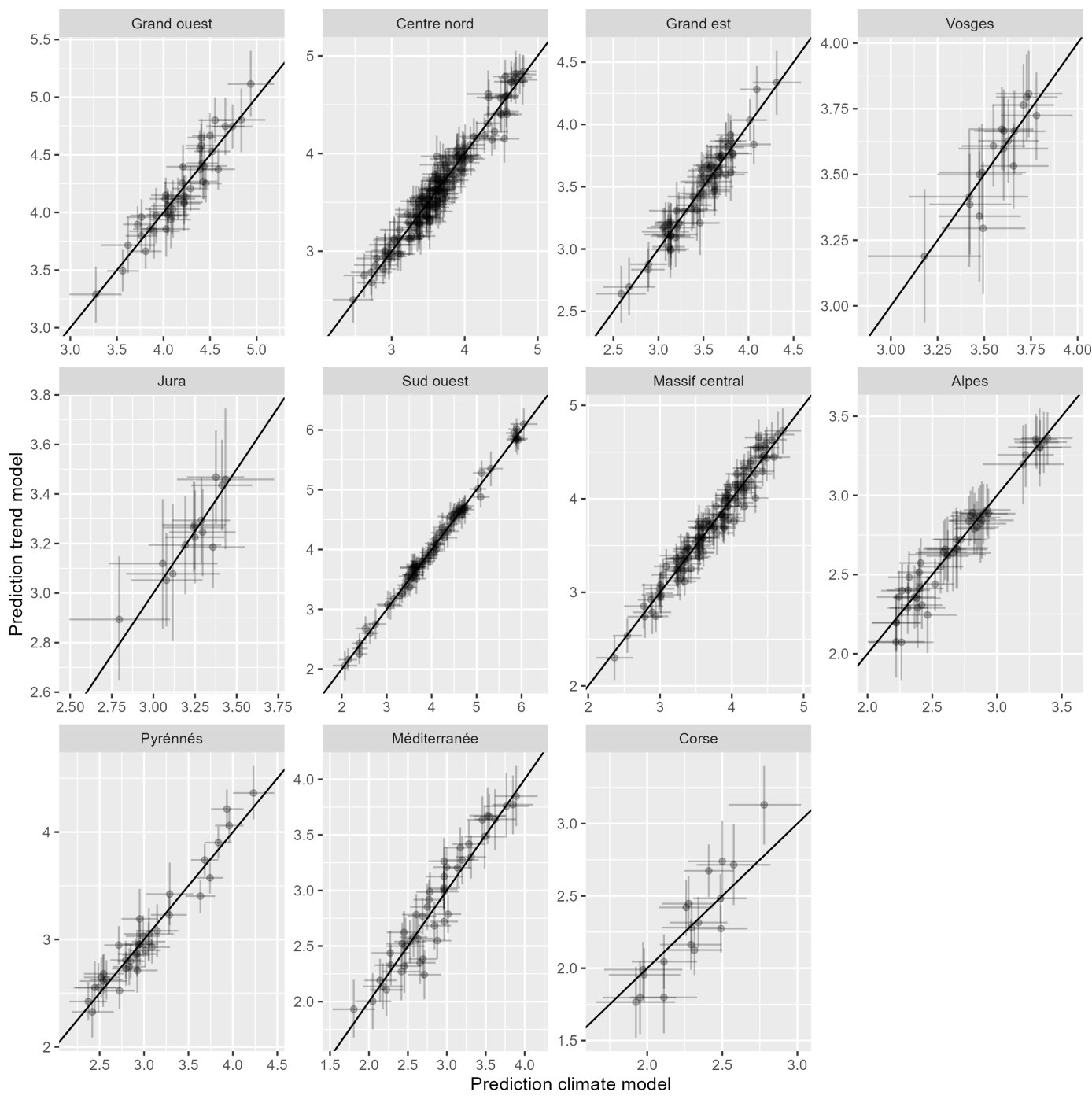
